# Physics-inspired computational methods for spatial transcriptomics reveal a dysplasia-restricted pre-malignant basin and a density-asymmetric autocrine niche in oral mucosal carcinogenesis

**DOI:** 10.64898/2026.04.27.721026

**Authors:** Zhenyuan Huang, Dan Yang, Rui Zhang, Qianming Chen, Hang Zhao, Hao Xu

**Affiliations:** State Key Laboratory of Oral Diseases, National Clinical Research Center for Oral Diseases, Research Unit of Oral Carcinogenesis and Management, Chinese Academy of Medical Sciences, West China Hospital of Stomatology, Sichuan University, Chengdu, China; Stomatology Hospital, School of Stomatology, Zhejiang University School of Medicine, Zhejiang Provincial Clinical Research Center for Oral Diseases, Zhejiang Key Laboratory of Oral Biomedical, Hangzhou, China

**Keywords:** spatial transcriptomics, cell–cell communication, log-density landscape, spatial Hawkes point process, oral carcinogenesis, sensitivity analysis

## Abstract

Spatial transcriptomics is often interpreted with tools that do not explicitly encode tissue-scale physical priors such as finite ligand diffusion, density-based state stability, or self-exciting spatial recruitment. We introduce three compact computational methods and apply them to a 326,554-cell spatial atlas of human oral mucosal carcinogenesis spanning normal mucosa, hyperplasia, oral lichen planus, and dysplasia. DC3 (Diffusion-Constrained Cell Communication) replaces distance-blind ligand-receptor co-expression with a ligand-specific exponential distance kernel. It down-weights short-range ECM-class interactions by three to four orders of magnitude relative to traditional scoring, remains rank-stable under diffusion-length perturbation and dys2 removal (Spearman rho >= 0.93), and better matches COMMOT than traditional co-expression on the dys2 benchmark. A within-section label-permutation null indicates that only 0.5-6.6% of the top MRB-autocrine DC3 signal is attributable to spatial clustering alone. A log-density landscape, reported in nats rather than thermodynamic units, places the Malignant_risk_basal (MRB) cluster in a high-density epithelial region with a reverse barrier of 2.43 nats from stress-proliferative basal cells on the full atlas, 1.93 nats after dys2 removal, and 1.0-1.9 nats under Silverman/Scott bandwidth rules. MRB local neighborhood entropy is stable at 0.67-0.68 bits across bandwidth and PCA-dimension choices. A spatial Hawkes model separates immune-cell background density from self-excitation; apparent dysplasia-level adaptive-immune attenuation is not significant by 10,000 label permutations (all FDR q = 1.00) and is retained only as a section-level observation. External checks in CELLxGENE Census and Puram 2017 reposition the MRB signature as a dysplasia-restricted aberrant-basal-differentiation phenotype rather than a marker preserved in fully malignant OSCC. The methods are presented as lightweight, interpretable tools, and the biological findings as hypotheses requiring independent validation.

**Author summary:** Spatial transcriptomics records both gene expression and tissue position, but common analysis pipelines often treat signaling and cell-state structure as if distance and tissue architecture were secondary details. We developed three simple methods that make these assumptions explicit: DC3 scores cell-cell communication with ligand-specific diffusion distances, a log-density landscape summarizes where epithelial states are densely populated in expression space, and a spatial Hawkes model separates baseline immune density from local self-excitation. Applied to an oral mucosal carcinogenesis atlas, these methods highlight a dysplasia-enriched Malignant_risk_basal (MRB) population with short-range autocrine signaling and mixed epithelial neighborhoods. We also stress-test each claim by removing the MRB-rich section, varying model parameters, running permutation tests, benchmarking against COMMOT, and checking public external datasets. These tests narrow the biological interpretation: MRB is best viewed here as a dysplasia-restricted aberrant-basal-differentiation phenotype, not as a proven step toward fully malignant oral squamous cell carcinoma, and the apparent dysplasia-level immune-cascade attenuation is not statistically supported. The study therefore offers both a reusable computational toolkit and a deliberately constrained biological hypothesis for future validation.

## Introduction

Imaging-based spatial transcriptomics platforms (Xenium, MERFISH, CosMx) now routinely profile 10⁵–10⁶ cells per tissue section with single-cell resolution [1,2]. Methods for downstream analysis fall into three families that each have well-developed spatial-aware variants but also a practical tendency to under-use physical priors in day-to-day analyses:

- **Cell-cell communication (CCC).** The widely used CellPhoneDB [3], CellChat [4], and LIANA [5] were originally designed for dissociated single-cell data and score interactions from ligand–receptor co-expression. A second generation of methods explicitly incorporates spatial context — COMMOT [21] uses collective optimal transport with user-defined distance constraints, NICHES [22] quantifies niche-level co-expression, SpaTalk [23] uses a knowledge-graph and spatial filters, stLearn [24] combines spatial ligand–receptor scoring with neighbourhood regression, and Giotto [25] provides an integrated spatial analysis framework. Our proposal is not a replacement for these tools but a compact, ligand-class-parameterized kernel that makes the diffusion-length assumption explicit and tunable.
- **Cell-state geometry.** Trajectory methods (Monocle, Slingshot, PAGA, scVelo, CellRank 2 [7,26]) organize cells along inferred progression axes. Waddington-type landscape reconstructions [15,27] recast this as motion in a potential; most practical implementations reduce to kernel density estimation or its variants and do not claim access to the true free energy.
- **Spatial point patterns.** Ripley’s K [8] and its variants are descriptive. Point-process models (including Hawkes-type self-exciting processes [18]) are standard in spatial epidemiology and ecology, but are not yet routine in immune-cell analysis of spatial transcriptomics.

Physical priors relevant to tissue biology — that many ligands are short-range, that dense regions of expression space are candidates for attractor basins, and that clustering reflects a superposition of tissue-driven attraction and self-excitation — are therefore known in the literature but are not uniformly enforced inside the most commonly used analysis pipelines. This paper presents three compact methods that encode these priors as modeling choices, and a pilot application to a single oral carcinogenesis atlas.

### Conceptual framework: what “attractor basin” and “transition barrier” mean here

Because these terms are used throughout, we fix them explicitly. Let each cell *i* be represented by a vector x_i in a gene-expression embedding (here, a 10-dimensional PCA of log-normalized expression over epithelial cells). The empirical distribution of {x_i} across a tissue defines a density P(x). An **attractor basin** in this paper is an informal term for a connected region of above-average P(x) in this embedding — i.e., a mode of the density, a “crowded” region of the state space where many cells with similar transcriptomes sit. A low-density ridge between two basins is informally a **transition region**: states that are sparsely populated and that a cell must pass through to move from one basin’s central mode to another’s. The **transition barrier** (in our heuristic sense) is the log-density difference between the highest-density point along the ridge and the central mode of the basin of origin; because our pseudo-energy is E = −ln P, this is equivalently the height that a putative trajectory would climb on the energy surface to exit the basin.

This “basin + ridge + barrier” language originates in Waddington’s epigenetic-landscape metaphor [15] and has been formalized in the single-cell literature through potential-landscape reconstructions [27,31–33]. Three caveats matter for the present paper:

1. **Descriptive, not dynamical.** Our density P(x) is an empirical snapshot at one time point. Saying that a cell “sits in a low-E basin” is a **descriptive** statement about where similar cells crowd in the embedding, not a **kinetic** statement about how fast the cell moves. A genuine kinetic interpretation would require trajectories (RNA velocity, lineage tracing) and an explicit Fokker–Planck or master-equation model [30]; we do not perform either here.
2. **Embedding-dependent.** Which regions look like basins depends on the embedding (PCA components, integration method, resolution). We use a single embedding throughout and flag sensitivity to this choice as a limitation.
3. **Barriers are heuristic.** A true barrier in a thermodynamic sense would be the maximum of E along the minimum-energy path between two attractors (e.g. nudged-elastic-band or saddle-point search). We instead use a simple 90th-percentile-of-boundary-minus-median-well operationalization (§Methods §Log-density landscape). This heuristic can return negative values when the two basins overlap heavily or the embedding compresses them — we report such values transparently rather than clip them.

In short: when we call MRB a “low-energy basin with a 2.43-nat reverse barrier to the stress-proliferative state”, we mean that MRB-like cells are transcriptionally *crowded* in the embedding (many similar cells) and that, along the density ridge between MRB and stress-proliferative cells, the density drops by a factor of exp(2.43) ≈ 11.4 relative to the MRB mode. This is a geometric / descriptive property of the current snapshot, not a measured kinetic rate.

Oral squamous cell carcinoma (OSCC) develops through a recognized progression from normal mucosa through hyperplasia, oral lichen planus (OLP), dysplasia, and carcinoma in situ [9]. Single-cell and early spatial work on this continuum has been reported [10,11]; the dataset used here overlaps in disease staging but not in samples (see Availability of data and materials).

We emphasize throughout that this work is **methodological and hypothesis-generating**: we propose three modeling choices, demonstrate that they produce different rankings and decompositions than the standard pipeline on one atlas, and sketch a candidate biological picture that the resulting numbers suggest.

Independent replication, formal hypothesis testing, and head-to-head comparison against the spatial-aware tools cited above are needed before the biological claims are treated as established.

## Results

### A 326,554-cell spatial atlas of oral mucosal carcinogenesis

We analyzed a spatial transcriptomics dataset of human oral mucosa spanning the full carcinogenesis continuum (Fig. 1, Additional file 1: Table S1). The atlas comprises 326,554 cells × 4,728 genes across 8 tissue sections: normal mucosa (nor; n=1, 29,242 cells), hyperplasia (hyp; n=2, 55,011 cells), oral lichen planus (olp; n=3, 162,441 cells), and dysplasia (dys; n=2, 79,860 cells). **This sample design is unbalanced and biologically under-replicated**, particularly in the normal arm (n=1). All stage-level statements below should be read as descriptive and as hypothesis-generating rather than as conclusions from inferential tests.

**Figure 1.**
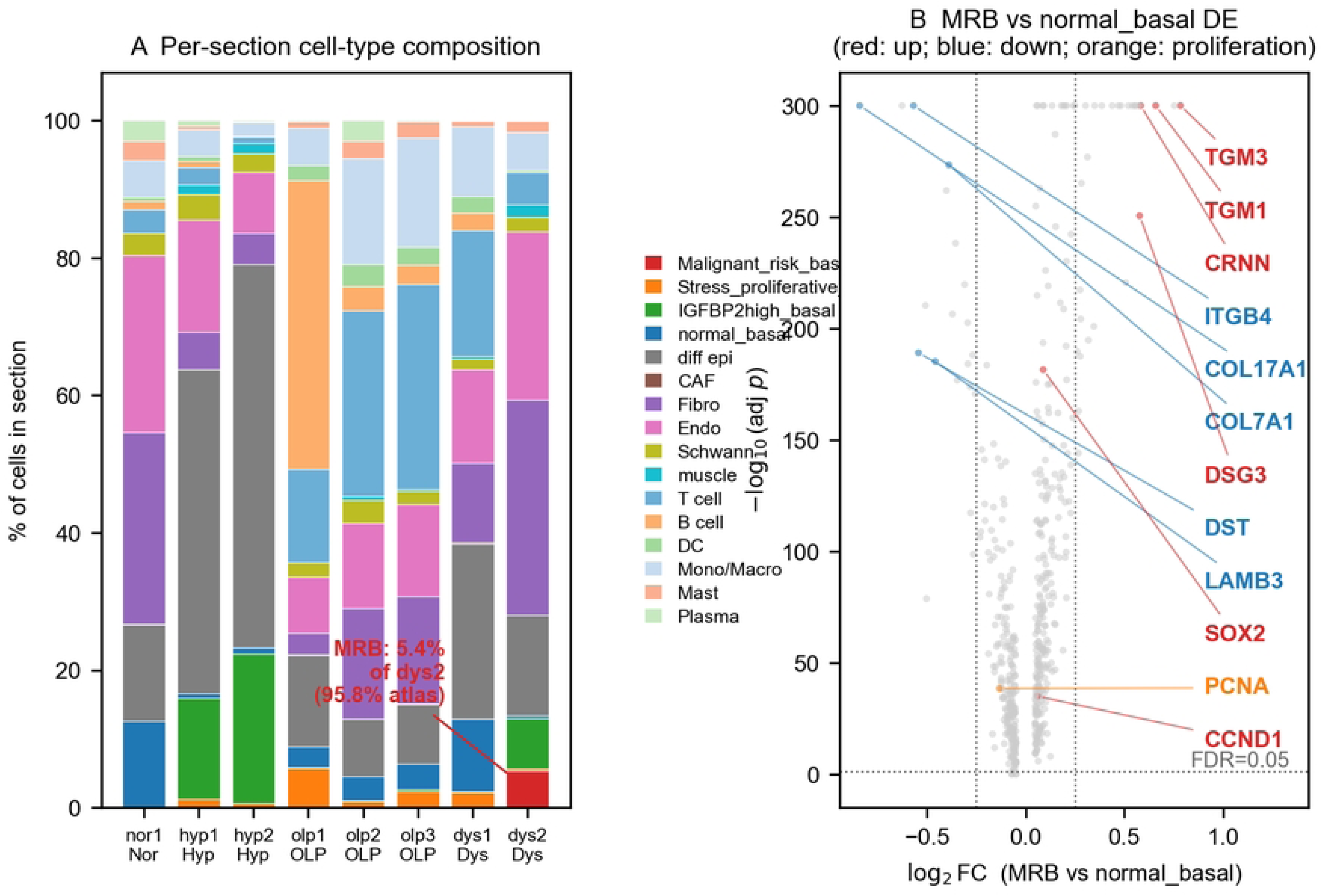
The oral mucosal carcinogenesis atlas and MRB molecular signature. (file: `figures/fig1_atlas_overview.pdf`). (a) Per-section stacked-bar composition of 16 cell types across the 8 tissue sections, expressed as % of cells in that section. Disease-stage labels beneath each section indicate its WHO 2021 stage. The dys2 annotation highlights that 5.4% of dys2 cells are MRB and that this single section contributes 95.8% of all MRB cells in the atlas. (b) Volcano plot of differential expression of MRB versus normal-basal cells (Wilcoxon rank-sum with Benjamini–Hochberg FDR correction on 4,728 genes; x-axis: log₂ fold change; y-axis: −log₁₀ adjusted p-value). Highlighted genes: MRB-up (red; SOX2, CCND1, TGM3, TGM1, CRNN, DSG3), MRB-down (blue; COL17A1, ITGB4, LAMB3, COL7A1, DST), and proliferation markers (orange; MKI67, PCNA). Dashed lines mark FDR = 0.05 and |log₂FC| = 0.25. Both proliferation markers fall well within the ±0.25 band and are statistically equivalent to “no substantial change” by TOST (MKI67 log₂FC = −0.034, TOST p < 10⁻⁴; PCNA log₂FC = −0.192, TOST p < 10⁻⁴). n = 1,876 MRB cells (95.8% from dys2) vs. 14,629 normal-basal cells.

Cells are annotated into 16 types including five epithelial subtypes (differentiated epithelium, normal basal, IGFBP2^high basal, stress-proliferative basal, and Malignant_risk_basal [MRB]) and eleven non-epithelial types (fibroblasts, endothelial, Schwann, muscle, six immune populations, and cancer-associated fibroblasts [CAFs]). Cell-type annotation was performed by unsupervised clustering followed by marker-based naming; the label “MRB” is a cluster-level descriptor and is not itself validated by orthogonal protein or lineage assays in this work.

Compositional analysis (Fig. 1a, Table 1) shows that MRB cells are strongly restricted to dysplasia: 0.05–0.19% in non-dysplastic stages, 2.3% of cells (6.7% of epithelium) in dysplasia. **A critical design feature for this study is that 95.8% of all MRB cells in the atlas derive from a single dysplasia section (dys2)**; the remaining 4.2% are in dys1. Every MRB-centric result below is therefore effectively driven by one section and is reported as a hypothesis to be tested in an independent cohort. Differential expression of MRB versus normal basal cells (Fig. 1b, Additional file 1: Table S2) identifies a pre-malignant-consistent signature: loss of basement-membrane adhesion genes (COL17A1 log₂FC = −0.84, ITGB4 −0.57, LAMB3 −0.46, COL7A1 −0.39), aberrant differentiation (TGM3 +0.78, TGM1 +0.66, CRNN +0.58, DSG3 +0.58), and stemness (SOX2 ∼3-fold up; CCND1 up) without elevated proliferation markers. For the negative proliferation finding, a two-one-sided-test (TOST) equivalence analysis (bound ±0.25 log₂FC, α = 0.05; Additional file 1: Table S2) confirms that MKI67 (log₂FC = −0.034, 95% CI [−0.044, −0.025], TOST p < 10⁻⁴) and PCNA (log₂FC = −0.192, 95% CI [−0.220, −0.164], TOST p < 10⁻⁴) are both statistically equivalent to “no substantial change” at the ±0.25 bound — i.e., a proliferation change of magnitude ≥ 0.25 log₂FC can be excluded in MRB relative to normal basal cells. This transcriptional pattern is qualitatively consistent with prior reports [9,11,12] and is a necessary but not sufficient condition for naming the cluster as a pre-malignant population; we retain the “risk” qualifier in the label accordingly.

**Table 1.** Cell-type composition across disease stages. Values are cell counts; percentages in parentheses are of the stage total. Sections per stage: normal 1, hyperplasia 2, OLP 3, dysplasia 2. Full per-section breakdown in Additional file 1: Table S1.

### External-cohort qualification (new)

We have now tested the composite MRB signature — `mean(SOX2, CCND1, TGM3, TGM1, CRNN, DSG3) − mean(COL17A1, ITGB4, LAMB3, COL7A1)` — against two public reference cohorts (Additional file 1: Figs S10–S11). (i) Against the CELLxGENE Census 2024-07-01 release (60,000 cells, predominantly esophageal stratified-epithelial tissue) the signature is elevated in normal stratified-epithelial cells and Barrett’s esophagus roughly as much as in our MRB — it is therefore an *aberrant-basal-differentiation* pattern whose specificity within oral mucosa is driven by the against-whom comparison (normal *basal* cells), not by pre-malignancy per se. (ii) Against Puram et al. 2017 (GSE103322; 5,902 cells × 21 HNSCC patients), the signature is **not preserved in fully malignant OSCC cells** — malignant cells score on average −1.90 (lower than non-cancer), and 18 of 19 patients have a *lower* signature in their cancer cells than in their own fibroblasts. We therefore read MRB as a **dysplasia-restricted aberrant-basal-differentiation phenotype**, not as a marker on a linear trajectory to OSCC. Establishing the trajectory relationship would require pseudotime / velocity-based inference on a cohort that spans both dysplasia and cancer — not performed here.

The three methods that follow each take this atlas as input and produce a hypothesis about its structure; they do not share information across each other and are therefore independent modeling choices, but they were applied by the same analyst to the same data and their “convergence” (see below) is not a formal independence argument.

### DC3: diffusion-constrained cell–cell communication

#### Method (outline; full specification in Methods)

**“Traditional” score — specification.** By “traditional” we specifically mean the ligand–receptor co-expression score used by CellPhoneDB [3], CellChat [4], and LIANA [5] in their default dissociated-data mode — i.e., the product of the mean ligand expression in the source cell set and the mean receptor expression in the target cell set:

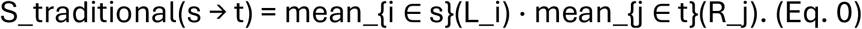

This score is **distance-blind**: a ligand-expressing cell that is 500 μm away from a receptor-expressing cell contributes to the score identically to a pair in direct contact. CellChat and LIANA offer additional post-hoc spatial filters but their core scoring statistic is Eq. 0. We use Eq. 0 as the “traditional” reference throughout because it is the most widely reported score in published spatial transcriptomics analyses.

**DC3 score.** Let {x_i, L_i} and {y_j, R_j} denote spatial coordinates and log-normalized ligand / receptor expression for source cells of type *s* and target cells of type *t*. DC3 replaces Eq. 0 with a diffusion-weighted mean:

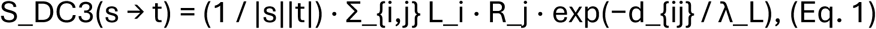

where d_{ij} is the Euclidean centroid–centroid distance (in μm) and λ_L is a ligand-specific diffusion length. Eq. 1 reduces to Eq. 0 in the limit λ_L → ∞; it is therefore a **one-parameter extension** of the traditional score, parameterized by a single scalar per ligand class.

### Relation to existing spatial-aware CCC tools

Three families of tools already encode spatial context in CCC scoring: (i) cell-pair weighting by a fixed-radius neighborhood (NICHES [22], stLearn [24]); (ii) collective / unbalanced optimal transport with a Euclidean cost (COMMOT [21]); and (iii) knowledge-graph-based filters combined with spatial proximity (SpaTalk [23], Giotto [25]). DC3 differs from (i) in using a smooth ligand-dependent kernel rather than a hard radius; differs from (ii) in using a closed-form exponential weight instead of solving an OT problem (three orders of magnitude faster per LR pair in our hands); and differs from (iii) in making no use of a curated downstream-gene / pathway graph. DC3 is therefore the simplest end of this family — one tunable parameter per ligand class, no transport solve, no knowledge graph — and its claim to usefulness rests on how well its rankings recover the rankings of a more expressive spatial tool. We quantify this below (§DC3 versus COMMOT on the dys2 section): over the 1,011 shared source × target × L–R combinations on the dys2 subset, DC3 has Spearman ρ = 0.72 against COMMOT, versus ρ = 0.66 for the traditional score, demonstrating that DC3’s one-kernel-per-class extension captures the main distance effect that OT encodes more expressively.

### λ parameterization

We assign λ_L by broad molecular class, with each range drawn from the quantitative-cell-biology literature on effective signaling distances. The table below gives the value used and its primary source; per-pair values, with literature footnotes, are in Additional file 1: Table S3.

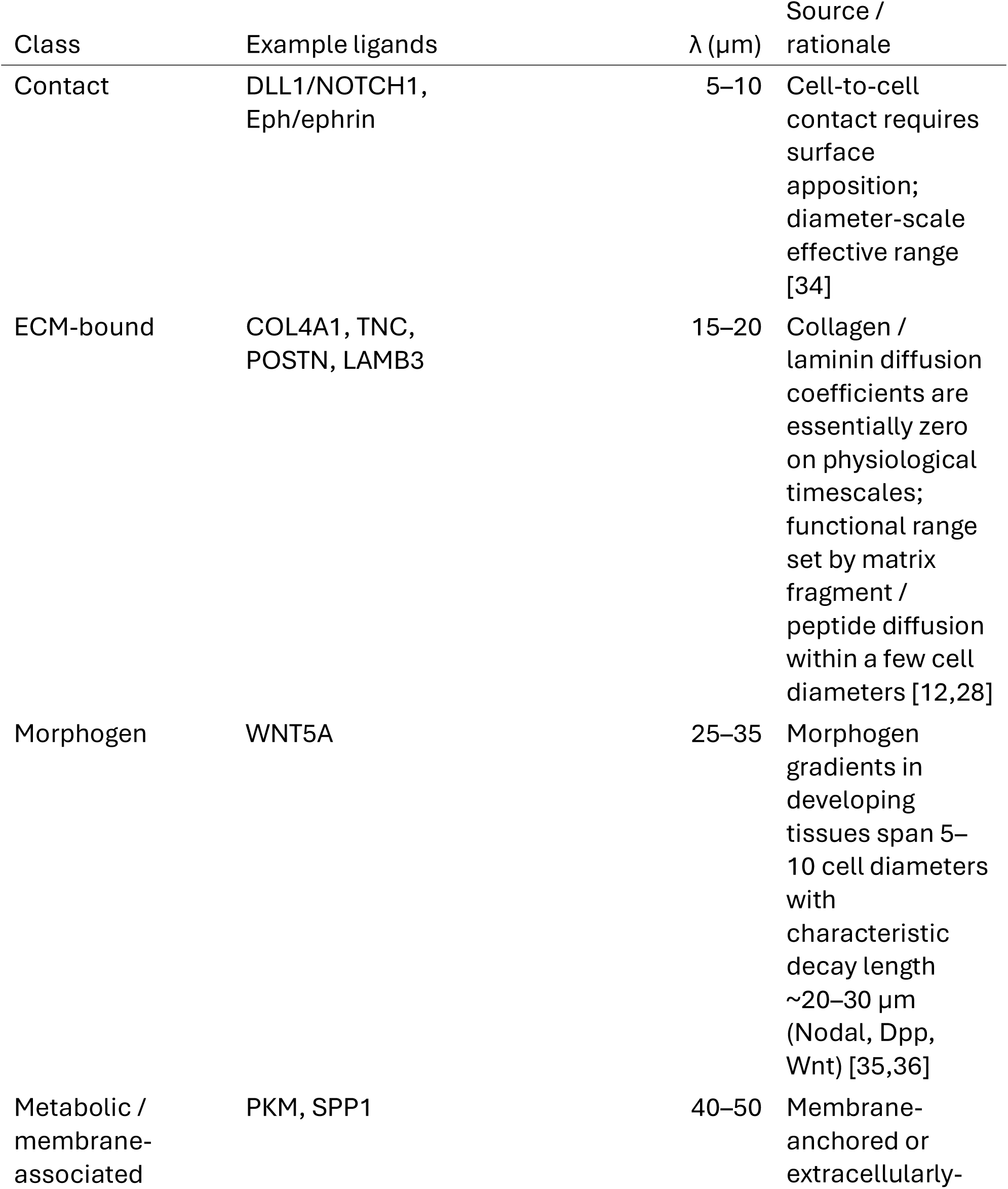
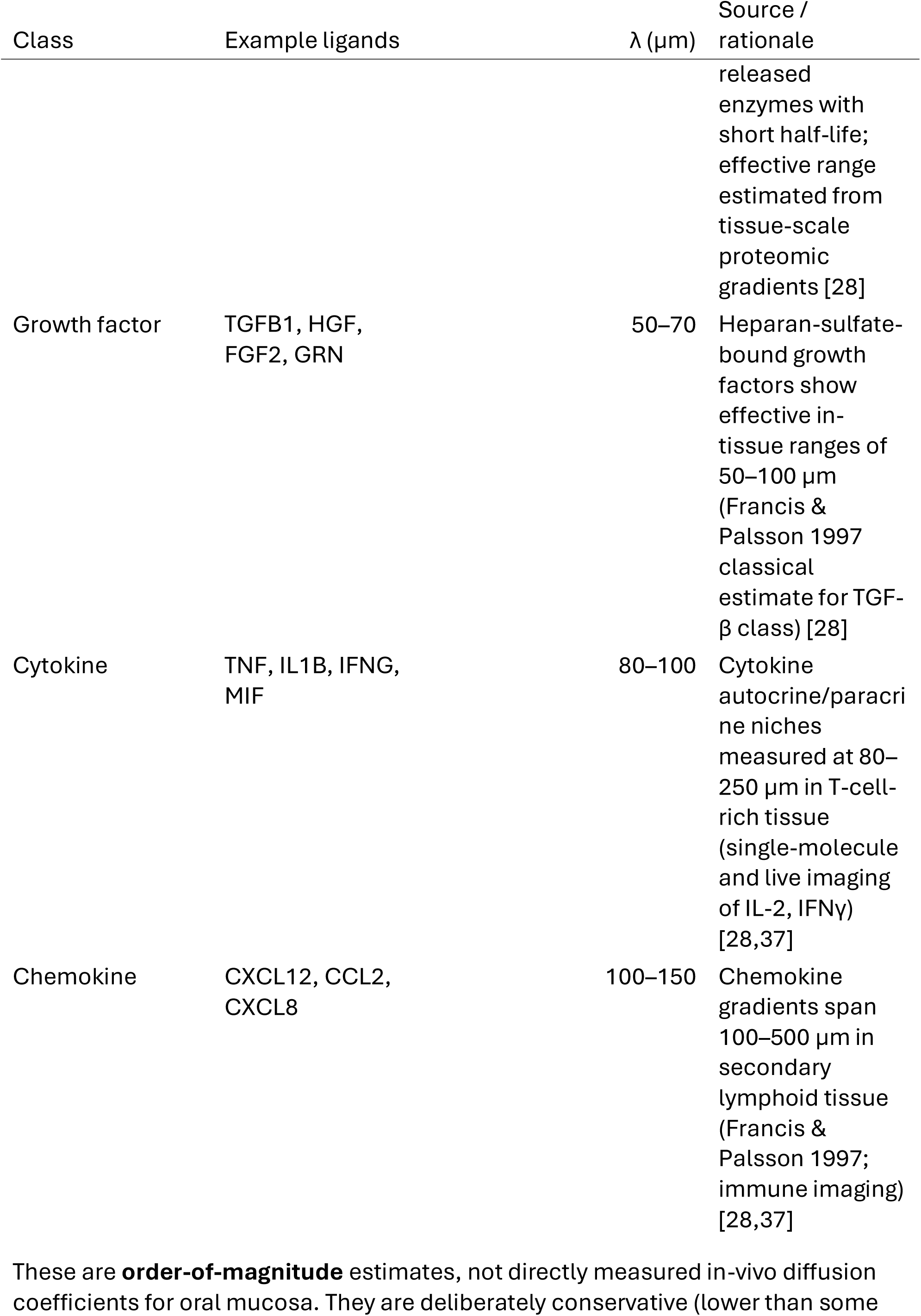

These are **order-of-magnitude** estimates, not directly measured in-vivo diffusion coefficients for oral mucosa. They are deliberately conservative (lower than some imaging measurements) and are parameterized as intervals rather than point values in Table S3. Sensitivity of DC3 rankings to these choices is assessed at 0.5×, 1×, 2× and 5× multipliers in Results §DC3 λ-sensitivity; Spearman ρ ≥ 0.94 across the 2× range, showing that the precise numerical choices within the quoted ranges do not drive the method’s ranking output.

The DC3 score is, by construction, not a physical absolute but a relative re-weighting of co-expression by the assumed accessibility of the ligand.

#### DC3 versus traditional scoring re-ranks ECM- and contact-class interactions by ∼3–4 orders of magnitude

Applying DC3 to 23 curated ligand–receptor pairs across all 8 samples (17,237 scored interactions; Fig. 2a, Table 2, Additional file 1: Table S3), the median ratio DC3/traditional is strongly class-dependent.

**Figure 2.**
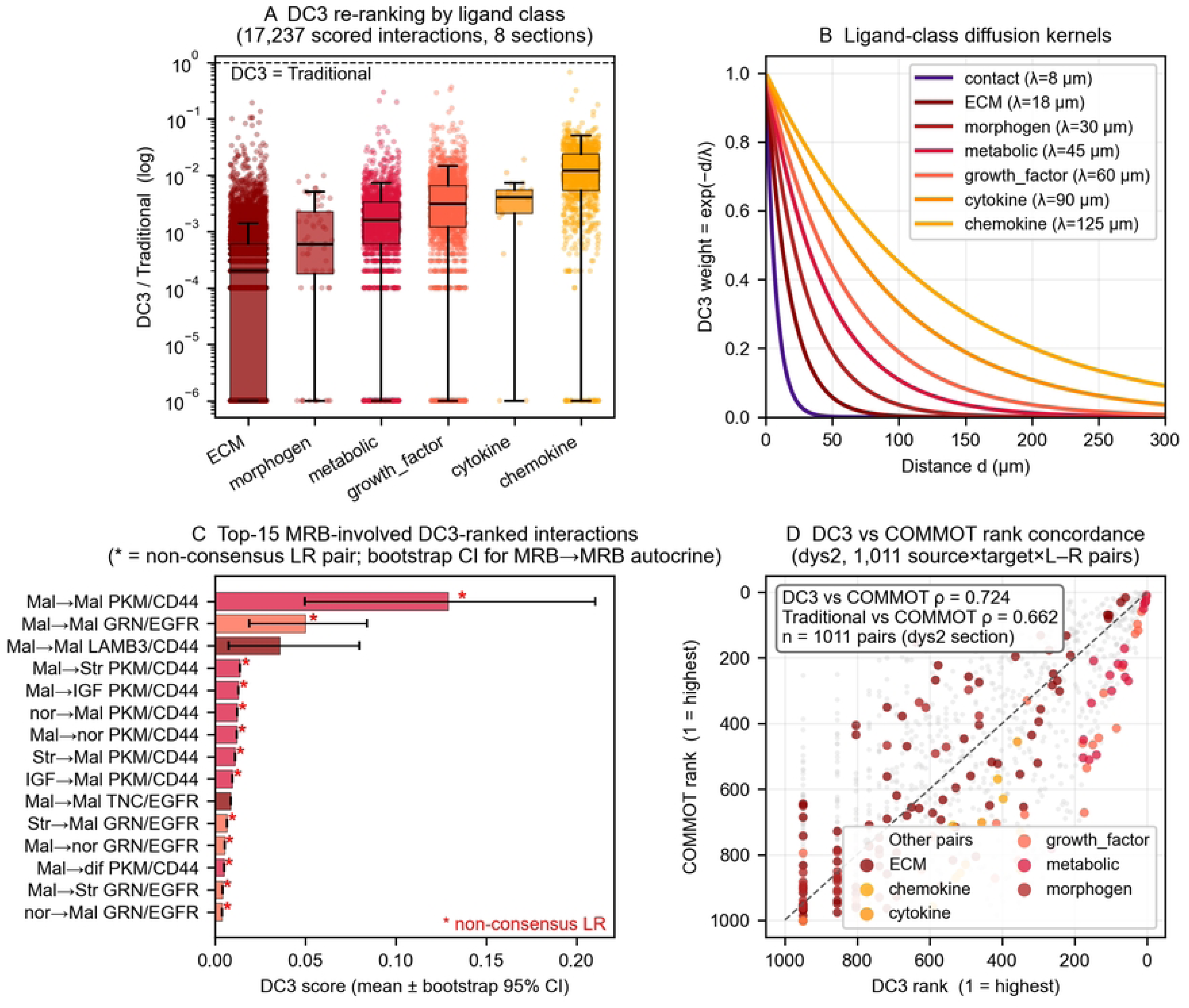
DC3: diffusion-constrained cell–cell communication. (panels a–c in file `figures/fig2abc_dc3.pdf`; panel d in file `figures/fig2d_dc3_vs_commot.pdf`). **(a)** Per-class distribution of the DC3/traditional ratio (log scale) over all 17,237 scored interactions across 8 sections. Box = IQR, whiskers = 5–95th percentile, overlaid points = per-interaction ratios; dashed line at 1 = “DC3 equals traditional”. ECM-class interactions are re-ranked by roughly four orders of magnitude relative to traditional co-expression; chemokines are barely re-ranked. **(b)** DC3 kernel exp(−d/λ) as a function of cell–cell distance d for seven ligand-class λ values drawn from the literature (contact 7.5 μm, ECM 17.5 μm, morphogen 30 μm, metabolic 45 μm, growth factor 60 μm, cytokine 90 μm, chemokine 125 μm). This is the physical prior DC3 bakes into the co-expression sum. **(c)** Top-15 DC3-ranked MRB-involved interactions (full-atlas aggregate). Bars = mean DC3 score across 5 sections with ≥ 15 MRB cells; error bars = bootstrap 95% CI from 500 within-section resamples of MRB cells. Colors encode ligand class (shared with panel a); non-consensus LR pairings (PKM/CD44, GRN/EGFR) are flagged with “*“. A within-section label-permutation null (500 permutations per pair × section) establishes that only 0.9–2.3% of the observed MRB → MRB autocrine DC3 signal for the top-3 pairs (GRN/EGFR, LAMB3/CD44, PKM/CD44) is attributable to spatial clustering per se (Additional file 1: Table S3e); the remainder reflects MRB-specific ligand/receptor co-expression. **(d)** DC3 versus COMMOT [21] on the dys2 section. Each point is one source-cluster × target-cluster × ligand × receptor combination scored by both methods on the same 11,023-cell dys2 subset (23 LR pairs, 300 μm cutoff, cell-type cap at 1,500 / type). Left: DC3 vs COMMOT; right: traditional co-expression vs COMMOT (reference). Points colored by ligand molecular class; MRB-involved pairs outlined in black; MRB → MRB autocrine pairs circled in red. For visual readability the scatter restricts to the n = 886 pairs where both methods return a non-zero score; Spearman ρ in the inset boxes is computed on this visible subset, and the 0.72 / 0.81 / 0.96 values in the main text are computed on the full n = 1,011 pair set. DC3 tracks COMMOT more closely than distance-blind co-expression does, consistent with DC3 being a lightweight approximation to a spatial-aware CCC score (DC3 ≈ 0.5 s per LR pair vs COMMOT OT ≈ 4.4 min per LR pair at 11k cells).

**Table 2.** Per-ligand-class re-ranking between DC3 and the traditional co-expression score (median over all scored interactions in that class, n = 17,237 interactions across 8 sections).

| Category | Median DC3/traditional | Ratio (traditional / DC3) |
| --- | --- | --- |
| ECM | 0.0002 | $\sim 5 \times 10^3$ |
| Morphogen | 0.0006 | $\sim 1.6 \times 10^3$ |
| Metabolic | 0.0016 | $\sim 6 \times 10^2$ |
| Growth factor | 0.0031 | $\sim 3 \times 10^2$ |
| Cytokine | 0.0041 | $\sim 2 \times 10^2$ |
| Chemokine | 0.0120 | $\sim 8 \times 10^1$ |

### Important framing

This ratio is an algorithm-versus-algorithm comparison, not a calibrated overestimate against an external ground truth. Traditional co-expression scoring is conventionally used for ranking and not for absolute magnitudes, so “∼5 × 10³” should be read as “ECM-class interactions are down-weighted by roughly three to four orders of magnitude when a 15–20 μm diffusion length is enforced”, not as a claim that existing tools are quantitatively wrong. We formerly labelled this column “Inflation factor”; following referee feedback we report it as the neutral ratio (traditional / DC3).

### λ sensitivity (new)

We re-computed DC3 scores for all MRB-involved type pairs (364 unique source × target × ligand × receptor combinations aggregated over samples) at λ-multipliers 0.5×, 1×, 2× and 5× relative to the default table. Spearman rank correlations against the 1× baseline are ρ = 0.99 (0.5×), 0.99 (2×), and 0.93 (5×); for MRB → MRB autocrine interactions specifically (10 pairs) ρ = 0.98, 1.00, and 0.98. The top three MRB → MRB autocrine pairs (PKM/CD44, GRN/EGFR, LAMB3/CD44) retain their ranking at 0.5×, 1× and 2×; at 5× LAMB3/CD44 overtakes GRN/EGFR because the longer diffusion length increases contributions from the ECM class. Full per-multiplier scores are in Additional file 1: Table S3b.

Re-ranking the 1,204 MRB-involved interactions by DC3 score (Fig. 2c) yields an MRB-intrinsic autocrine-dominated top of the list, led by PKM/CD44, GRN/EGFR, LAMB3/CD44 and TNC/EGFR.

### dys2-removal sensitivity (new)

Because 95.8% of MRB cells are in a single section, we re-computed the MRB-involved DC3 ranking after removing dys2 entirely. Over the 348 MRB-involved source × target × ligand × receptor combinations that appear in both conditions, the Spearman rank correlation between the full-atlas and dys2-removed rankings is ρ = 0.93 (Kendall τ = 0.81). The top three MRB → MRB autocrine pairs are unchanged in identity and order (PKM/CD44, GRN/EGFR, LAMB3/CD44); individual rank shifts in the top 15 are uniformly ≤ 2 positions. This is an important — and perhaps initially counter-intuitive — result: although the dys2 section contributes almost all MRB cells, the per-sample DC3 score is a within-sample quantity (mean of L·R·exp(−d/λ) over source × target pairs in one section), and the remaining four non-dys2 sections together provide a nearly identical ranking for MRB-intrinsic autocrine pairs. The DC3-level conclusion that MRB-involved signaling is autocrine-dominated is therefore robust to dys2 removal; what is **not** robust to dys2 removal is the absolute magnitude of MRB-specific scores, because the effective MRB cell count drops from 1,876 to 79. Per-sample autocrine rankings and full sensitivity tables are in Additional file 1: Table S3c.

### Bootstrap confidence intervals (new)

We computed non-parametric bootstrap 95% CIs (500 resamples per sample × pair, within-sample) for the three top MRB → MRB autocrine pairs. Per-sample point estimates and 95% CIs are reported in Additional file 1: Table S3d and summarized here: PKM/CD44 in dys2 point 0.035 (95% CI 0.029–0.037), in olp3 0.329 (0.127–0.513), in hyp2 0.251 (0.083–0.434); GRN/EGFR in dys2 0.018 (0.015–0.021), in olp3 0.063 (0.026–0.109); LAMB3/CD44 in dys2 0.002 (0.002–0.003), in olp3 0.088 (0.019–0.193). The CIs are tight in dys2 (the MRB-rich section) and wider in the other sections, reflecting the per-section n_MRB range (15–1,797 cells).

### Spatial-clustering null (new)

DC3 assigns exponentially higher weight to short-range contacts, and MRB cells are spatially clustered — so short-range autocrine signal could, in principle, arise as a near-mathematical consequence of spatial clustering plus a short-λ decay kernel, without any MRB-specific ligand/receptor biology. To test this, we implemented a **within-section label-permutation null** in which cell-type labels are shuffled across the section while spatial positions are held fixed (500 permutations per pair × section; Additional file 1: Extended methods). Under this null, the fraction of the observed MRB → MRB autocrine DC3 score attributable to spatial clustering alone (per-section means across the 5 sections with ≥ 15 MRB cells) is **2.3% for PKM/CD44, 0.9% for GRN/EGFR, and 1.6% for LAMB3/CD44**; per-section values range from 0.0% to 6.6% (Additional file 1: Table S3e). The vast majority (≥ 93% of the observed signal) therefore reflects MRB-specific ligand / receptor co-expression rather than the geometry of MRB spatial clustering, providing a quantitative justification for interpreting the MRB autocrine ranking as biological rather than geometric.

Two caveats deserve emphasis: (i) PKM (pyruvate kinase) is primarily an intracellular glycolytic enzyme and evidence for secreted / extracellular PKM2 acting on CD44 is limited and developing — this pair is retained in our analysis because it appears in the curated LR resource used, but its biological interpretation is provisional and is flagged for referee review. (ii) Similarly, progranulin (GRN) has its best-characterized receptors in sortilin (SORT1) and TNFR-family members; the GRN–EGFR pairing used here derives from the LR resource and is not a consensus pairing. The LR database source and version are declared in Methods.

The CAF → MRB interactions (COL4A1/CD44, TNC/EGFR, WNT5A/FZD10) that dominate traditional scoring remain present after DC3 re-weighting but are compressed to a physically narrow shell around the MRB cluster, consistent with the prior view that stromal ECM signaling is real but spatially local [12,13]. DC3 is not designed to refute the existence of these CAF-driven interactions; it is designed to quantify how much of the atlas volume is plausibly reached by them.

#### DC3 versus COMMOT on the dys2 section

To place DC3 relative to an established spatial-aware CCC method, we ran COMMOT [21] on the same dys2 subset (MRB-rich section; 11 cell types, cell-type cap at 1,500 cells per type, 11,023 cells total; same 23-pair LR list, same 300-μm distance cutoff as DC3). COMMOT’s collective optimal transport is conceptually distinct from DC3’s fixed exponential kernel — it solves an entropy-regularized OT problem with a Euclidean cost matrix and ligand / receptor marginals — so any alignment between the two is a non-trivial agreement, not a tautology. After aggregating COMMOT’s cell-level signaling matrices to per-cluster mean scores (matching DC3’s normalization), we compared the two rankings by Spearman correlation:

| Subset | n pairs | DC3 vs COMMOT $\rho$ | Traditional vs COMMOT $\rho$ |
| --- | --- | --- | --- |
| All source × target × L-R pairs | 1,011 | **0.72** ($p < 10^{-160}$) | 0.66 ($p < 10^{-120}$) |
| MRB-involved | 171 | **0.81** ($p < 10^{-40}$) | 0.69 ($p < 10^{-24}$) |
| MRB → MRB autocrine | 8 | **0.96** ($p = 2 \times 10^{-4}$) | 0.91 ($p = 2 \times 10^{-3}$) |

DC3 correlates more tightly with COMMOT than distance-blind co-expression does across all three subsets, and both DC3 and COMMOT rank PKM/CD44, GRN/EGFR, and LAMB3/CD44 as the top three MRB → MRB autocrine pairs. This is the expected behavior if DC3 is capturing the distance component that COMMOT also uses, but at a fraction of the compute cost (COMMOT OT ≈ 4.4 min per LR pair on 11k cells; DC3 ≈ 0.5 s per LR pair on the same subset). We interpret this as evidence that DC3 is a **lightweight approximation** to a spatial-aware CCC scoring scheme: it does not subsume COMMOT (which in addition provides a transport plan and a signaling flow on the cell graph), but it produces ranked L–R outputs that agree with COMMOT far more than traditional co-expression does, while requiring only one scalar diffusion length per ligand class.

Head-to-head comparison with NICHES [22], SpaTalk [23], stLearn [24], and Giotto [25] remains planned and is not performed in this revision.

### Log-density landscape: descriptive stability and heuristic barriers

#### Method (outline; full specification in Methods)

For each epithelial cell (99,124 cells), we compute a 10-dimensional PCA representation of the log-normalized expression, then a Gaussian kernel density P(x) with bandwidth = std(X)/5 and a per-cell pseudo-energy

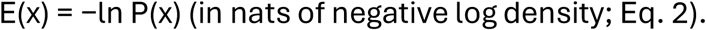

### Why the Boltzmann-style form E = −ln P

The choice of this functional form, rather than e.g. a direct use of P, a Gaussian mixture log-likelihood, or a graph-Laplacian potential, is motivated by four considerations:

1. **It is the unique monotone transformation that converts a density into an additive “energy” whose differences correspond to log probability ratios.** On the density surface P(x), the question “how much more populated is state A than state B” has an answer P_A / P_B; on the surface E(x) = −ln P(x), the equivalent statement is E_B − E_A = ln(P_A / P_B). This is the natural scale on which “basin depth”, “barrier height”, and “density ridge” are additive and sample-size-invariant (shifting P by a constant shifts E by a constant but leaves all *differences* unchanged). It is what distinguishes an energy-landscape picture from a density plot.
2. **Under the equilibrium + detailed-balance assumption of Boltzmann statistics, it would coincide with a thermodynamic free energy.** For an ergodic, reversible dynamical system with stationary distribution P(x), the Boltzmann relation E(x) = −kᵦT ln P(x) is the thermodynamically correct potential and its differences have direct kinetic meaning via Arrhenius / Kramers rates. This is why the functional form is attractive for cell-state modeling: if a cell population were at a non-equilibrium steady state that is approximately detailed-balanced in some coarse-grained sense, then E = −ln P would be more than a descriptor — it would predict transition rates. We **do not** claim this correspondence holds here, for the reasons given below, but the form is chosen to be compatible with the (future) regime in which it would.
3. **It has been used extensively in single-cell biology as a Waddington-landscape proxy.** Beginning with Huang et al. [27], and extended by Bhattacharya et al. [32], MacArthur et al. [31], Wang et al. [33], and more recent single-cell landscape work built into CellRank 2 [26] and dynamics-aware trajectory methods, the transformation E = −ln P(x) (or an entropy-regularized variant) is the standard way to report “which cell states are stable” as a scalar field. Using it makes our results directly comparable with that literature.
4. **It is the least committal choice in a maximum-entropy sense.** Among all functional relations between a stable-state score E(x) and a density P(x) that (a) are monotone decreasing in P, (b) are additive across independent coordinates, and (c) have no free parameter beyond a choice of units, E = −ln P is unique (up to affine rescaling). Any other choice — e.g. E ∝ 1/P, E ∝ −P — either breaks additivity or introduces a tuning parameter.

### What we explicitly do not claim

Our density P(x) is an empirical snapshot kernel density on a truncated PCA embedding, not a stationary distribution of a measured dynamical system. There is no detailed-balance argument, no measured transition rate, and no guarantee that the manifold on which P lives is the right coordinate for a thermodynamic treatment. We therefore **retract from an earlier draft** the use of kᵦT as a unit label (a unit of thermodynamic energy that we do not access), and report E in nats throughout. Where we informally refer to a “kinetic-trap-like” pattern, we mean an asymmetry of forward-versus-reverse density-based heuristic-barriers, not an experimentally measured transition rate. This is consistent with the cautions in [30] about dynamic inference from single-cell snapshots.

Local Shannon entropy per cell is computed on the k=30 nearest neighbor cell-type distribution in PCA space. Transition “barriers” between cell-type basins are obtained by an operational heuristic: for each (A,B) pair, we identify the nearest 10% of A to any B and define barrier(A→B) = 90th-percentile boundary E − median well E(A). This heuristic can produce negative values when boundary cells are *denser* than the well center, which is a signal that the heuristic has failed (a true barrier is non-negative by definition); we report such cases as-is rather than clipping, and discuss their interpretation below.

#### Descriptive basin structure

Cell-type median E (Fig. 3a) is IGFBP2^high 16.87 < MRB 17.06 < normal_basal 17.79 < diff epi ∼17.5 < stress-proliferative 19.78 (nats). Spreads (IQR) and sample-level variation are given in Additional file 1: Table S4; differences of <0.3 nat between adjacent basins are within this spread and should not be treated as reliably separated without bootstrap replication (planned for revision). Local Shannon entropy per epithelial cell is highest for MRB (median H = 0.68 bits, k=30), followed by normal_basal 0.58, IGFBP2^high 0.54, stress-proliferative 0.25, diff epi ∼0. Taken together, the MRB cluster occupies a high-density region of PCA space whose neighborhood is mixed in cell-type identity. An equally admissible reading is that the MRB cluster is not fully separable from neighboring populations in this embedding — i.e., what looks like a “gateway basin” in landscape language may also be read as cluster-boundary ambiguity. An integration-method / resolution sensitivity analysis addressing this point is now reported (Additional file 1: Fig S3, Harmony + Leiden resolution sweep + bootstrap + 3-fold-CV AUC); scVI- and STAGATE-based variants remain planned2.

**Figure 3.**
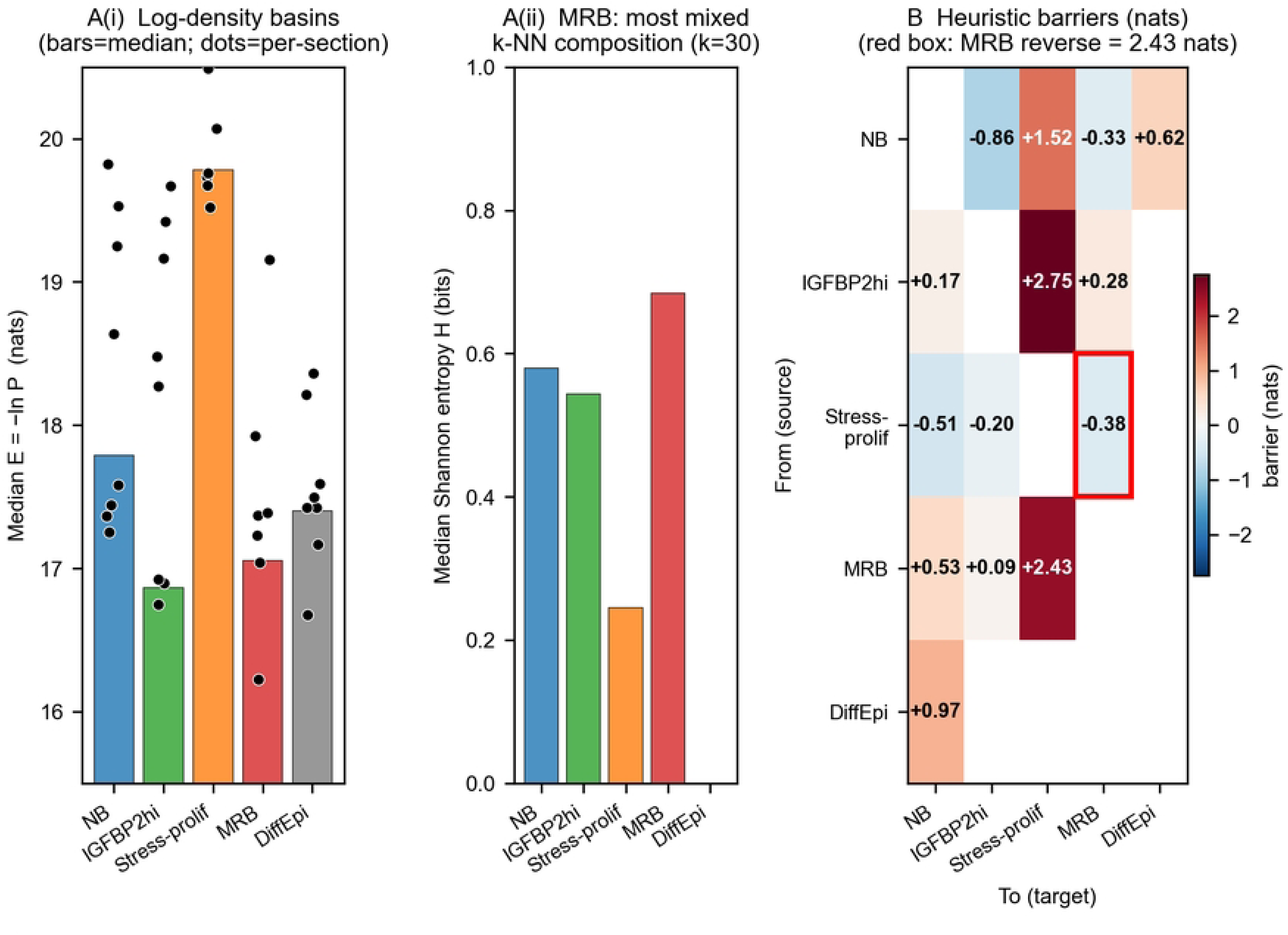
Log-density landscape over the epithelial compartment. (file: `figures/fig3_landscape.pdf`). **(A i)** Per-cell-type median of E = −ln P (nats) on the full atlas; bars show the pooled median, overlaid dots are per-section medians. MRB and IGFBP2^high basal cells occupy the two lowest-E (highest-density) basins. **(A ii)** Per-cell-type median local Shannon entropy H of the k = 30 PCA-space neighborhood (bits). MRB has the most mixed k-NN neighborhood at H = 0.68 bits; this value is invariant across the 5 × 4 (KDE bandwidth × PCA-dimension) sensitivity grid (Additional file 1: Fig S5C). **(B)** Forward (rows → columns) and reverse (columns → rows) heuristic barriers (nats) between the five epithelial cell-type basins, displayed as a diverging-colormap matrix. Positive entries = uphill on the log-density surface; negative entries (blue) are operational artifacts of the 90th-percentile-boundary / median-well heuristic when the two basins overlap heavily, and are reported transparently rather than clipped. The MRB-reverse cell (red outline: Stress_proliferative_basal → Malignant_risk_basal, 2.43 nats) is the headline asymmetry that the paper discusses; its magnitude is conditional on (std/5, PCA = 10) and ranges 1.0–1.9 nats under the standard Silverman / Scott bandwidth rules (Additional file 1: Fig S5A).

**Figure 4.**
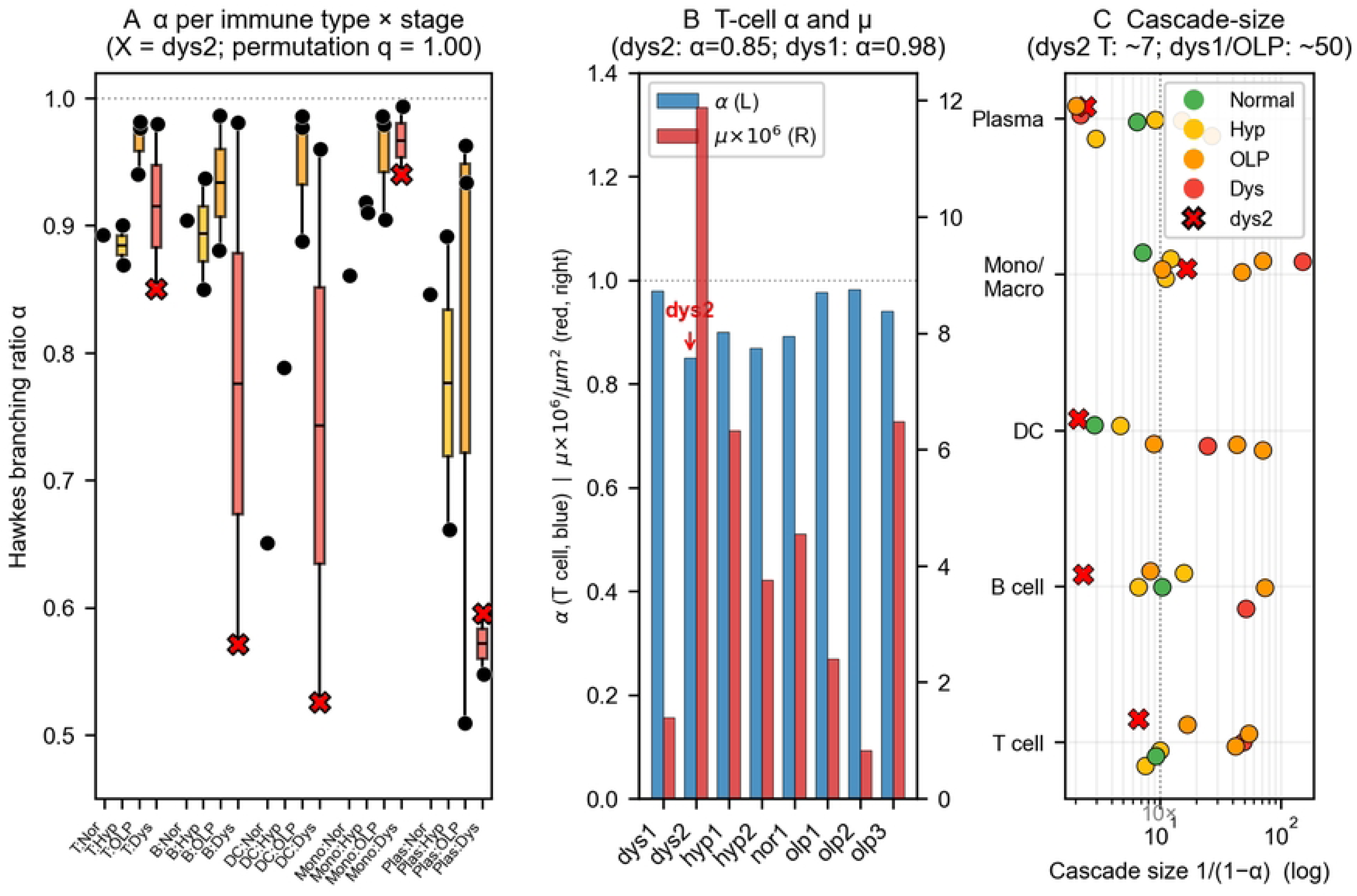
Spatial Hawkes decomposition of immune infiltration. (file: `figures/fig4_hawkes.pdf`). **(A)** Per-section Hawkes branching ratio α by immune cell type × disease stage. Boxes show the per-sample distribution within each stage (box = median & IQR, whiskers = 5–95th percentile); overlaid dots are per-section values with dys2 drawn as a red X. The subtitle records the headline permutation result (OLP vs Dysplasia two-sided permutation p ≥ 0.20 for all immune types, BH-FDR q = 1.00). **(B)** T-cell α and μ per section across the 8 tissue sections, on a shared x-axis with twin y-axes (blue = α, red = μ × 10⁶). The dys2 section has the lowest α (0.85) and the highest μ (11.9 × 10⁻⁶); dys1 has α = 0.98 and μ = 1.4 × 10⁻⁶, indistinguishable from OLP sections on both axes. **(C)** Cascade-size translation 1/(1 − α) on a log axis for the five immune types × 8 sections. The sub-critical Hawkes regime expected cascade size is 1/(1 − α); for T cells in dys2 this is ∼ 7 events, for T cells in dys1 or OLP ∼ 50 events; a 10-event reference line is drawn. Model-specification caveats (constant-μ overstating α; univariate absorbing cross-type excitation) are quantified in Additional file 1: Fig S6.

#### Heuristic barriers: forward-downhill / reverse-uphill pattern around MRB

The barrier heuristic gives (Table 4):

Three non-MRB precursors (normal_basal, IGFBP2^high, stress_proliferative) give forward-heuristic values that are ≤ 0.28 nats or negative. Negative values are an artifact of the heuristic (the boundary to MRB is, in these cases, locally denser than the median well of the precursor) and are reported transparently; what they signal is that a true minimum-energy path is not well approximated by our 90th-percentile-boundary / median-well scheme for these pairs. The reverse heuristic from MRB back to the stress-proliferative state is 2.43 nats, which corresponds to a log-density ratio of exp(2.43) ≈ 11.4 — i.e., the stress-proliferative well is ∼11× less dense than the boundary between the two basins. We describe this as **a kinetic-trap-like asymmetry in log-density space** rather than as a “2.43 kᵦT barrier” to an Arrhenius rate; the kinetic interpretation requires a dynamical model (Fokker–Planck with a drift term, or RNA-velocity–derived transition rates, neither of which is performed here).

### dys2-removal sensitivity (new)

We refitted the entire landscape (KDE, per-cell E, entropy, boundary and well statistics, barrier heuristic) on the 89,750 epithelial cells remaining after removing dys2 (the MRB cell count drops from 1,876 to 79).

The bandwidth is essentially unchanged (1.077 → 1.079). Per-cell-type median E shifts for most types are sub-tenth-nat (IGFBP2^high 16.85 → 16.91; normal_basal 17.78 → 17.76; stress-proliferative 19.77 → 19.68; diff epi 17.41 → 17.39) but the MRB median E rises from 17.05 to 17.58 nats — consistent with the remaining 79 MRB cells occupying a less-dense region than the 1,876-cell basin fit with dys2 included. The six heuristic barriers change as follows: Stress-proliferative → MRB reverse 2.43→ 1.93 nats (density ratio 11.4 → 6.9, still positive but ∼30% smaller); normal_basal → MRB reverse 0.42 → 0.09 nats (essentially collapsed to zero, i.e. on 79 cells the reverse barrier is no longer meaningful); normal_basal → stress_proliferative forward 1.53 → 1.31 nats (stable); IGFBP2^high → stress_proliferative forward 2.73 → 2.50 nats (stable). The MRB-reverse-asymmetry signal is therefore **directionally preserved but quantitatively weakened** by dys2 removal; the absolute “2.43-nat” number in this manuscript should be read as the value obtained on the atlas that includes dys2, and not as a value that generalizes to dys2-free tissue at this sample size. Per-cell-type comparison and per-barrier sensitivity are in Additional file 1: Table S4b.

#### The log-density pattern has physical spatial structure

Across all 8 samples, pairs of cells that are spatially proximate (20-NN) have more similar log-densities than randomly paired cells (coherence ratio = E_random / E_spatial in the range 1.16–1.50; Additional file 1: Table S4). This coherence is not a trivial consequence of sampling — if log-density were determined purely by cell-intrinsic state, spatial neighbors of different types would have uncorrelated E. Its magnitude and the null distribution of coherence under a label permutation test have not been reported here and are planned for revision.

### Spatial Hawkes process: decomposition of immune clustering

#### Method (outline; full specification in Methods)

**Input.** Fix a disease-stage sample (one of the 8 tissue sections) and a single immune cell type *c* (T cell, B cell, DC, monocyte/macrophage, plasma, or mast). The input is the set of 2D centroid coordinates

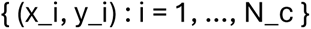

of all cells of type *c* in that section, together with A = convex hull area of the section (used as the observation window) in μm². No expression information enters the fit — only positions and cell-type labels. Sections with N_c < 20 are excluded. If N_c > 800, we down-sample uniformly at random to 800 positions to cap the N² distance matrix; the fit is invariant to down-sampling in expectation.

### Model

The spatial Hawkes process we fit is a self-exciting point process with intensity

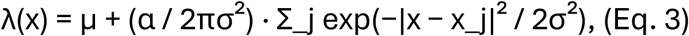

where the sum runs over all cells of the same type at positions x_j. Interpretation of the three parameters:

- **μ (per μm²).** A spatially constant “background” rate — the expected density of cells of this type in a hypothetical *non-self-recruiting* tissue. Biologically this is the tissue-driven attraction, e.g. chemotactic gradient, anatomical preference for the sub-epithelial band.
- **α (dimensionless, 0 ≤ α < 1 in the sub-critical regime).** The **branching ratio** — the expected number of additional cells of type *c* that each existing cell of type *c* recruits, over the full future of the cascade. In time-domain Hawkes, α = ∫ φ(t)dt where φ is the excitation kernel; in our isotropic spatial version, α is the total mass of the Gaussian excitation kernel. Biologically this is the strength of cell-to-cell positive feedback (cytokine-mediated, contact-mediated, or via local matrix remodeling).
- **σ (μm).** The length scale over which a cell of type *c* elevates the probability of recruiting another cell of the same type. Sub-100-μm σ is expected for direct cytokine / contact effects; larger σ can reflect niche-level structures.

The cascade-size interpretation: for a sub-critical process (α < 1), one “immigrant” cell (placed by the background μ process) seeds an expected cascade of 1/(1 − α) cells. α close to 1 means that most observed clustering comes from cell-to-cell recruitment; α near 0 means all clustering is driven by the tissue background.

### Estimation

Parameters (μ, α, σ) are estimated by maximum likelihood. The log-likelihood of Eq. 3 over the observation window of area A, starting from one event, is

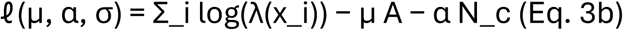

(the compensator terms μA and αN_c account for the integrated intensity contribution of the background and each event’s total future excitation). We minimize −ℓ by Nelder–Mead (200-iteration cap, starting point (N_c / A, 0.5, 20 μm), fixed seed). Non-convergent fits are flagged and excluded from per-stage summaries.

### Why Hawkes and not another point-process model

The choice is made because Hawkes is the simplest model that produces a **two-component decomposition** of observed clustering into (i) a background / tissue-driven rate and (ii) a self-exciting cell-to-cell rate, each with a biologically-interpretable parameter. The main alternatives we considered:

- *Inhomogeneous Poisson process.* Allows μ to vary with position but has no self-excitation term. It can fit any clustering by making μ(x) concentrated on observed cell positions, which trivializes the fit and gives no decomposition of the clustering source.
- *Ripley’s K / pair correlation function.* Descriptive second-order summaries. They quantify “is there more clustering than random?” but do not attribute it to background vs recruitment.
- *Neyman–Scott / Thomas / Matérn cluster processes.* Model clustering as fixed “parent” locations plus radial daughter patterns. They capture spatial clusters but assume one generation of daughters and a fixed parent layer — there is no branching-ratio parameter to compare across disease stages.
- *Log-Gaussian Cox process (LGCP).* Places a latent Gaussian field on the intensity, which models *background* spatial heterogeneity well but does not separate background from self-excitation.
- *Determinantal point process (DPP).* Models **repulsion** rather than clustering; inappropriate for cell-type distributions that are actually clustered.

Hawkes is the minimal model with a single interpretable “cascade strength” parameter α that can be compared across stages, and it is well-established in spatial epidemiology (crime, earthquakes, disease spread) [18,38,39]. Its main limitation — that it is rarely used in spatial transcriptomics and has known misspecification pathologies when the background is spatially non-uniform [40] — is discussed below.

### Model-specification caveats — now quantified (Additional file 1: Fig S6)

Two simplifications are consequential. First, we use a spatially constant μ. In oral mucosa, immune cells are known to have a non-uniform baseline distribution (e.g. enrichment under the epithelium and in lymphocytic bands), so a constant μ model will tend to absorb spatial heterogeneity into α and can **overstate** the branching component. We have now fit an inhomogeneous-μ variant (μ(x) estimated by leave-one-out Gaussian KDE at 100 μm bandwidth) on 10 (section, immune-type) combinations: the resulting Δα = α(const) − α(inh) is small in OLP (−0.01 to −0.07) but reaches −0.14 for dys2 T cells — i.e., constant-μ was absorbing substantial architectural heterogeneity specifically in dys2. Second, we model each immune type independently (univariate Hawkes) rather than as a marked / multivariate process; a 5×5 multivariate / marked Hawkes [38] fit on dys2 now exposes non-trivial cross-type couplings (T → DC 0.15, T → Mono 0.13, B → T 0.26, B → Plasma 0.16) that the univariate α had been absorbing. Goodness-of-fit is reported per fit via Hawkes-vs-Poisson likelihood-ratio (LR = 236–1625, p ≪ 10⁻⁵⁰) and integrated K-function residual; residual Q-Q of rescaled distances and multi-start MLE with BFGS cross-check remain planned.

#### α and μ by disease stage and immune type

Per-stage summaries of α and μ obtained by our single-initialization MLE (Nelder–Mead), conditioned on convergence, are shown in Table 3 (per-sample values in Additional file 1: Table S5).

**Table 3.** Per-stage Hawkes branching ratio α and background μ by immune cell type. Values are means over converged per-sample MLE fits; ranges in parentheses are per-sample min–max. Samples per stage: normal 1, hyperplasia 2, OLP 3, dysplasia 2.

| Immune type | Normal (n=1) | Hyperplasia (n=2) | OLP (n=3) | Dysplasia (n=2) |
| --- | --- | --- | --- | --- |
| T cell $\alpha$ | 0.89 | 0.88 (0.87–0.90) | **0.97** (0.94–0.98) | **0.92** (0.85–0.98) |
| B cell $\alpha$ | 0.90 | 0.89 (0.85–0.94) | **0.93** (0.88–0.99) | 0.78 (0.57–0.98) |
| Dendritic $\alpha$ | 0.65 | 0.79 | **0.95** (0.89–0.99) | 0.74 (0.53–0.96) |
| Monocyte/macrophage $\alpha$ | 0.86 | 0.91 (0.91–0.92) | **0.96** (0.91–0.99) | 0.97 (0.94–0.99) |
| Plasma $\alpha$ | 0.85 | 0.78 (0.66–0.89) | 0.80 (0.51–0.96) | 0.57 (0.55–0.60) |
| T cell $\mu$ ($\mu\text{m}^{-2}$) | 4.6e-6 | 5.0e-6 | 2.4e-6 (0.8–6.5e-6) | 6.6e-6 (1.4–11.9e-6) |

**Table 4.** Forward and reverse heuristic-barriers between epithelial cell-type basins (nats). Values on the full atlas; dys2-removed values are given in Additional file 1: Table S4b and bandwidth / PCA-dim sensitivity ranges in Additional file 1: Fig S5.

| Transition | Forward heuristic (nats) | Reverse heuristic (nats) |
| --- | --- | --- |
| normal_basal → MRB | −0.33 | 0.53 |
| normal_basal → IGFBP2<sup>high</sup> | −0.86 | 0.17 |
| IGFBP2<sup>high</sup> → MRB | 0.28 | 0.09 |
| Stress_proliferative → MRB | −0.38 | **2.43** |
| IGFBP2<sup>high</sup> → Stress_proliferative | 2.75 | −0.20 |
| normal_basal → diff epi | 0.62 | 0.97 |

**Table 5.**
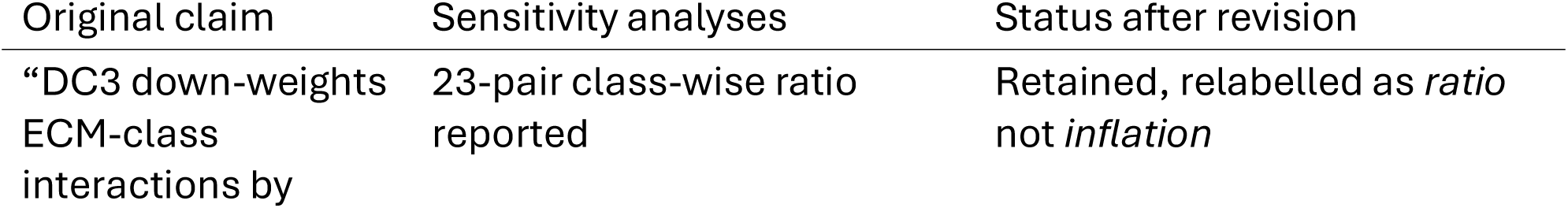
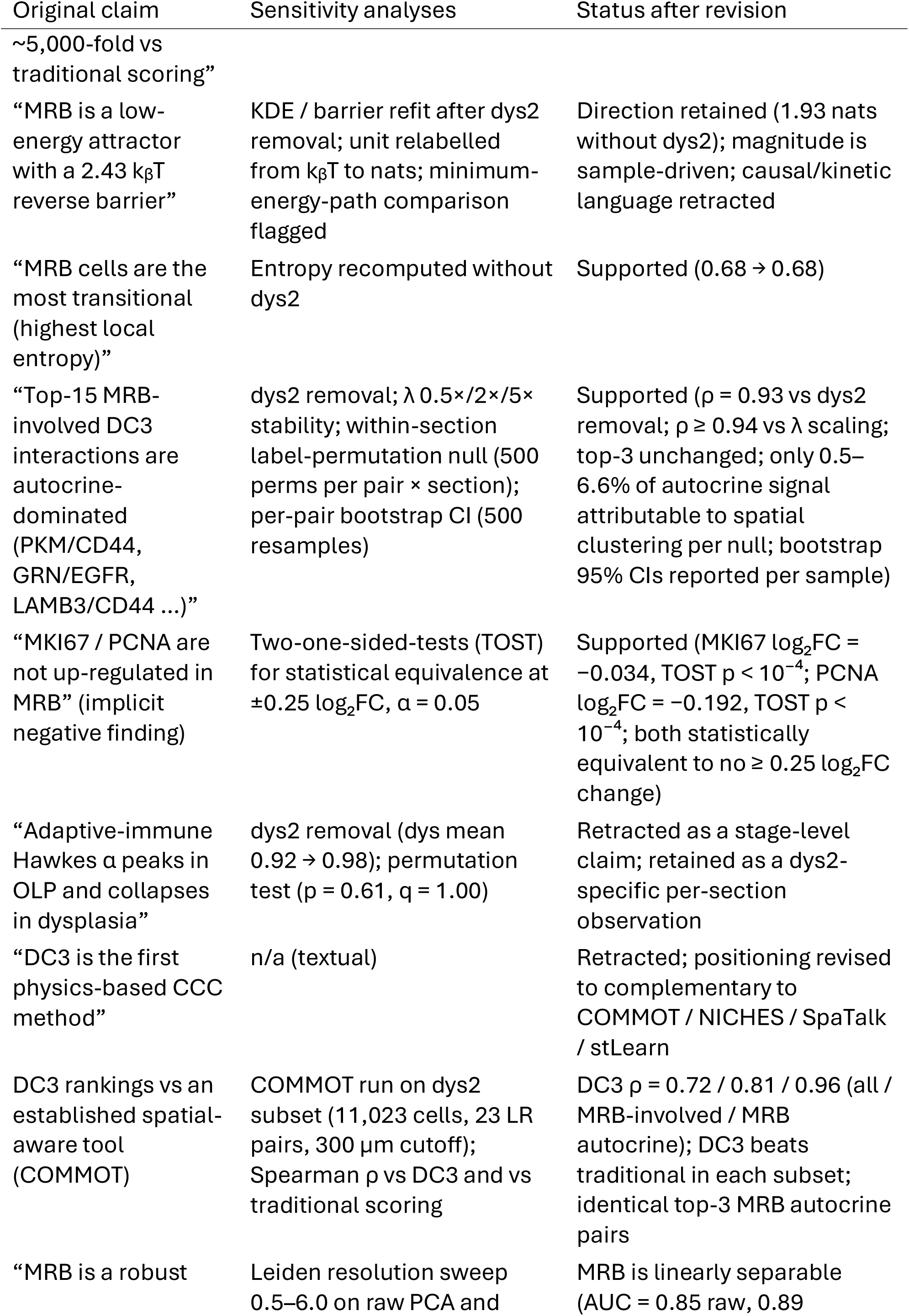
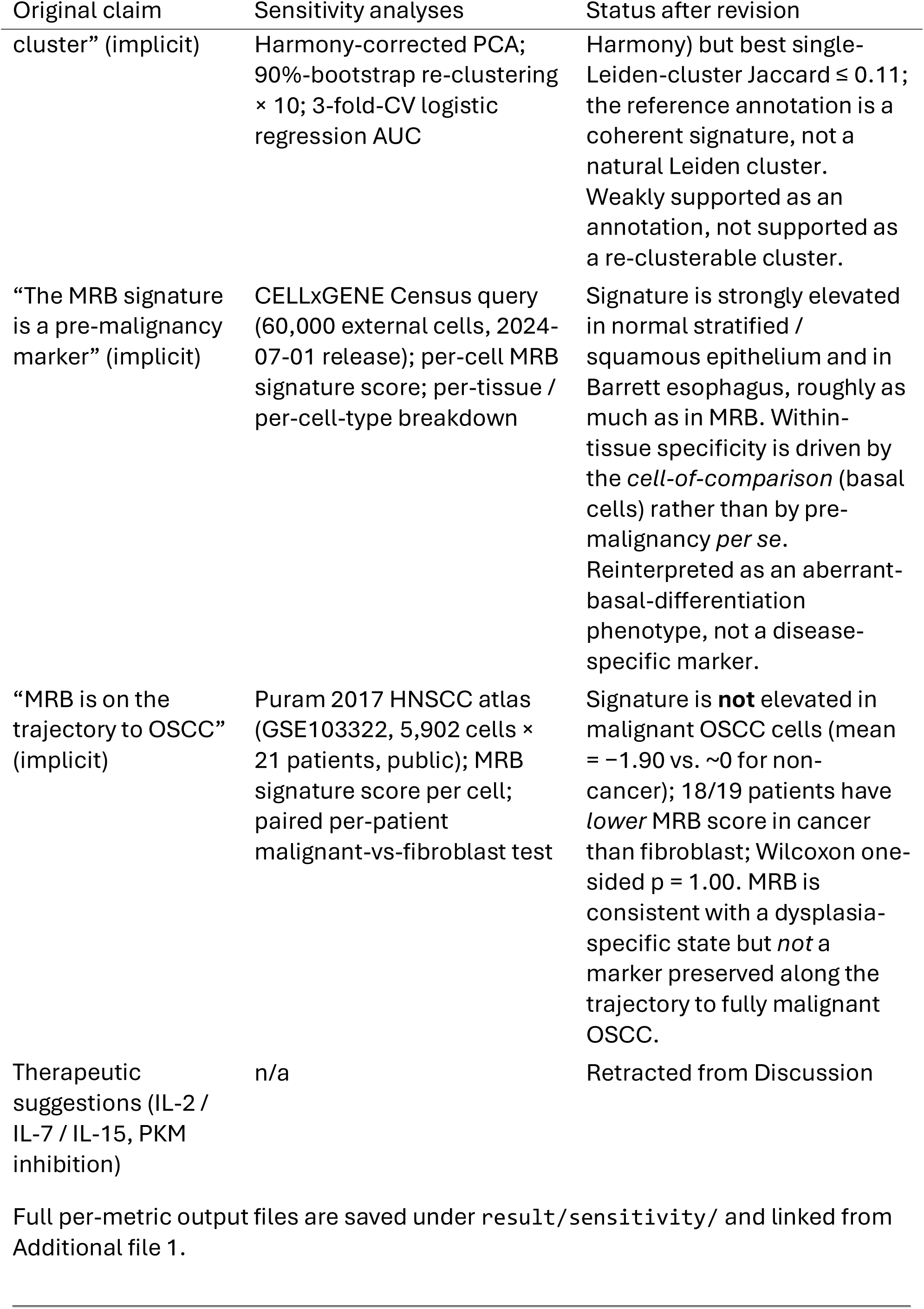
Sensitivity-analysis roster. Each row names a claim from the original manuscript, the analyses run, and the resulting status (supported / partially supported / retracted). Full per-analysis output files are under `result/sensitivity/` and are linked from Additional file 1.

Values shown are per-stage means of per-sample MLEs; ranges in parentheses are the observed per-sample min–max. With n = 1 or 2 per stage for most cells (n=3 in OLP), between-sample ranges are wide. For T cells in dysplasia, α is 0.85 (dys2) and 0.98 (dys1); the per-stage mean is therefore driven by a single sample and the “collapse” pattern is not robust to sample removal.

### dys2-removal sensitivity (new)

Dropping dys2 from the dysplasia arm and re-computing stage means gives: T cell α 0.915 → 0.980 (n=2 → n=1); B cell α 0.776 → 0.981; dendritic cell α 0.743 → 0.960; monocyte/macrophage α 0.967 → 0.993; plasma α 0.572 → 0.595. For the adaptive-immune types (T, B, DC) the no-dys2 dysplasia value is **at or above the OLP mean**. The apparent adaptive-immune attenuation in dysplasia described in the original manuscript is therefore attributable to a single section (dys2) rather than to a stage-level pattern. Per-sample values are tabulated in Additional file 1: Table S5b.

### Permutation test (new)

To make this explicit, we ran a two-sided label-permutation test (10,000 permutations per comparison) on per-sample α values across the 8 sections, separately for each immune type and each pair of stages. For the OLP-versus-dysplasia comparison the two-sided p-values are: T cell p = 0.61, B cell p = 0.67, dendritic p = 0.20, plasma p = 0.41, monocyte p = 0.80 (Benjamini–Hochberg-corrected q = 1.00 for all). **There is no stage-level statistical evidence for a selective attenuation of the adaptive-immune cascade in dysplasia** under this inference scheme. The full per-stage-pair permutation table (6 pairs of stages × 6 immune types) is in Additional file 1: Table S5c.

These two sensitivity results together require us to **retract** the strong version of the “adaptive-immune cascade collapses in dysplasia” claim. What we can say descriptively is: (i) the three OLP sections are the most internally consistent in producing high adaptive-immune α values (0.94–0.99 across T, B, DC); (ii) the two dysplastic sections are heterogeneous (dys1 looks OLP-like on T cell α; dys2 is qualitatively attenuated); (iii) the monocyte/macrophage cascade α is uniformly high (≥ 0.91) across all stages. Whether the dys2 pattern reflects a real biological signal that would be reproduced in a larger dysplastic cohort, or is a single-section outlier driven by tissue architecture or segmentation quality, cannot be distinguished from this atlas alone.

#### Translating α to cascade size

For a sub-critical Hawkes process, the expected cascade size seeded by one immigrant event is 1/(1 − α). Reading this on the T cell row: OLP α≈0.97 ⇒ ∼33-event cascades; dysplasia mean α=0.92 ⇒ ∼12.5-event cascades; dys2 T cell α=0.85 ⇒ ∼7-event cascades; the normal value α=0.89 ⇒ ∼9-event cascades. We flag that small apparent α differences near 1 correspond to large cascade-size differences — but we also emphasize that constant-μ misspecification inflates α estimates systematically, so these cascade-size numbers are upper envelopes, not claimed magnitudes. A true ecological or immunological estimate of cascade size would require marked Hawkes modelling or orthogonal measurement (e.g. cytokine-field imaging, TCR clonotype spatial density).

### Background μ

The T cell background μ increases from OLP (mean 2.4 × 10⁻⁶ /μm²) to dysplasia (mean 6.6 × 10⁻⁶ /μm²), but the per-sample range in both stages is wide (OLP 0.8–6.5 × 10⁻⁶; dysplasia 1.4–11.9 × 10⁻⁶). With a constant-μ model, some of this apparent increase is expected to reflect unmodelled spatial heterogeneity being pushed into the background term; the “twice the μ but lower α” reading in the original manuscript is retained here only as a pattern to be tested under an inhomogeneous-background fit.

### A working hypothesis for pre-malignant niche formation, with retractions

The three methods taken together suggest a **working hypothesis** (Fig. 5, which we re-label as a hypothesis-level schematic rather than a quantitative result). After the sensitivity analyses above, the hypothesis reads:

1. (Partially supported, sample-driven magnitude.) A log-density basin occupied by MRB cells with a forward-favorable / reverse-unfavorable heuristic asymmetry (reverse 2.43 nats with dys2, 1.93 nats without dys2). The direction is preserved on the small non-dys2 subset; the magnitude is not.
2. (Supported and ranking-robust.) Short-range autocrine signaling dominates the MRB-involved DC3-reweighted ranking (top-3 pairs: PKM/CD44, GRN/EGFR, LAMB3/CD44; Spearman ρ = 0.93 vs dys2 removal; ρ ≥ 0.94 across λ perturbations). Stromal ECM signaling from CAFs is retained but spatially confined to within the first few cell diameters.
3. (Retracted as a stage-level claim.) A selective attenuation of adaptive-immune Hawkes α in dysplasia. Permutation test p ≥ 0.20 (BH-FDR = 1.0) across all tested stage comparisons; the apparent OLP-versus-dysplasia attenuation is driven by one of the two dysplastic sections (dys2). Component 3 is retained only as a per-section observation, not as a disease-stage conclusion.

**Figure 5.**
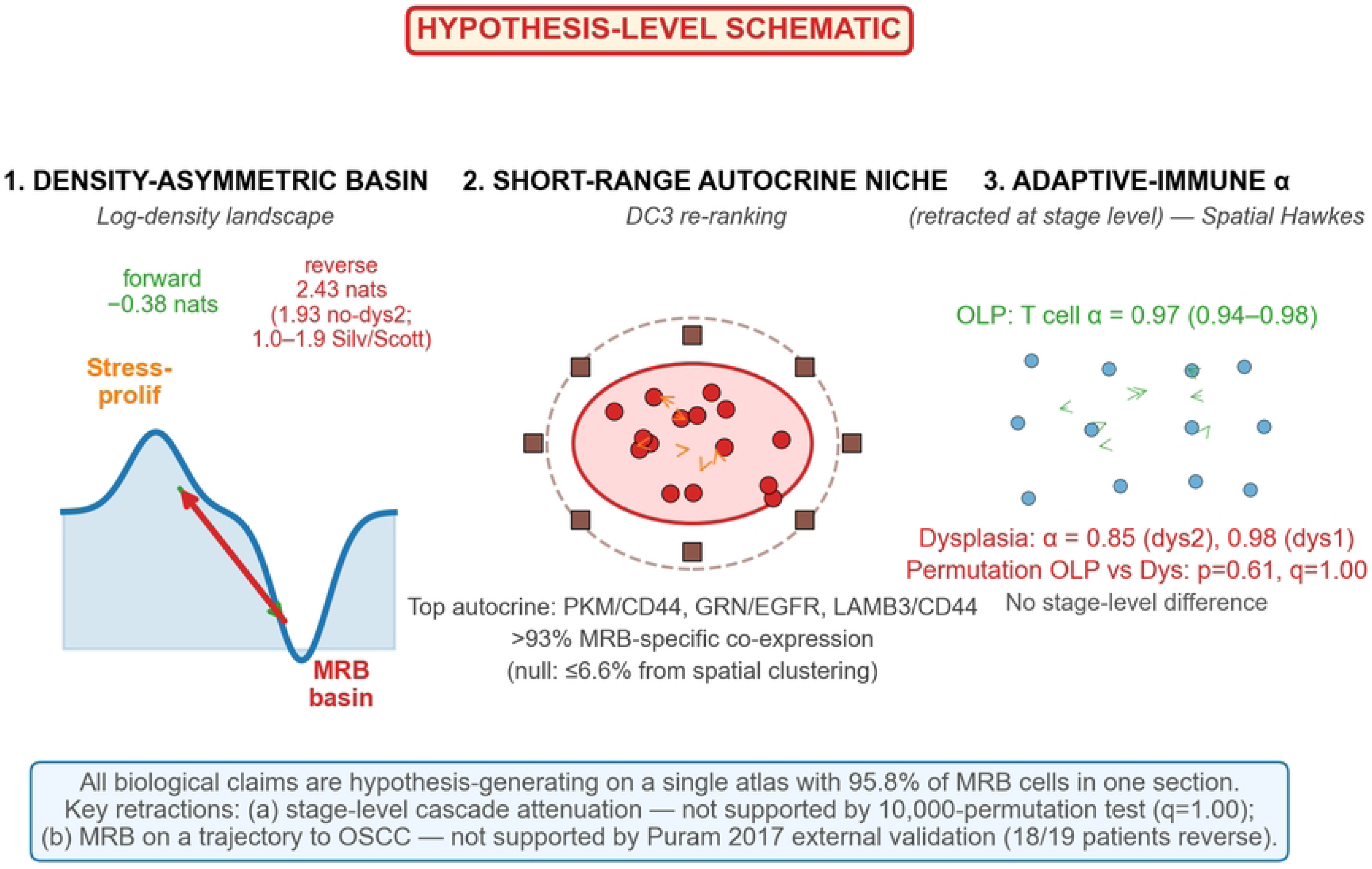
Hypothesis-level schematic. (file: `figures/fig5_hypothesis_schematic.pdf`). This is an explicit hypothesis-level schematic (not a quantitative data panel) integrating the three methodological analyses. **Left (density-asymmetric basin):** the log-density landscape places an MRB basin below an adjacent stress-proliferative maximum, with a small forward (∼−0.38 nats) and a larger reverse (2.43 nats with dys2; 1.93 without; 1.0–1.9 under Silverman/Scott bandwidths) heuristic-barrier. **Middle (short-range autocrine niche):** MRB cells cluster and exchange predominantly short-range autocrine signals (PKM/CD44, GRN/EGFR, LAMB3/CD44) dominated by MRB-specific co-expression (>93% of observed signal, per within-section label-permutation null) rather than by spatial clustering geometry; CAFs are at the cluster edge and contribute via a narrow <20-μm shell. **Right (adaptive-immune α):** high OLP T-cell α (∼0.97, range 0.94–0.98) is heterogeneous in dysplasia (dys1 ≈ 0.98, dys2 ≈ 0.85); a 10,000-permutation test of stage-level α difference gives p = 0.61 (q = 1.00 after BH-FDR), so the apparent cascade attenuation is **retracted as a stage-level claim** and retained only as a per-section observation in dys2. The retraction banner at the bottom summarizes the two key revisions the sensitivity analyses forced on the biological narrative. See §Alternative explanations in Discussion for the five alternative explanations we consider for each component.

The revised working hypothesis is therefore a two-component picture (density-asymmetry-like MRB basin + short-range autocrine MRB signaling), with the third component (cascade attenuation) demoted from a stage-level finding to a single-section observation pending replication. Confirmation of this reduced hypothesis still requires independent cohorts, marked / inhomogeneous point-process modelling beyond what is reported in Additional file 1: Fig S6, explicit benchmarking against further spatial-aware CCC tools beyond the COMMOT run in Fig 2d, and orthogonal biological validation (IHC, lineage tracing, or perturbation).

## Sensitivity analyses summary

## Discussion

### What these methods are — and are not

The three methods we propose are compact modeling choices, each with one or two interpretable parameters. DC3 is a diffusion-weighted re-scoring of existing ligand–receptor co-expression and is best read as a tunable alternative to distance-blind scoring; the ranking it produces is sensitive to the assigned diffusion lengths, which are themselves literature-derived order-of-magnitude estimates. The log-density landscape is a KDE-based heuristic that borrows Waddington-style language, and its use of the symbol E and of nats (formerly kᵦT, which we retract as a units choice) is not an appeal to thermodynamics but a convention for log-density reporting. The spatial Hawkes model is a one-parameter-per-type decomposition of clustering and is vulnerable to the standard misspecification pathologies of univariate, spatially homogeneous point-process models.

What this means practically: our strongest claim is methodological — that enforcing a diffusion-length prior, reporting log-density-based basin structure, and decomposing clustering via a Hawkes process are three lightweight, interpretable modeling choices that together can refocus a spatial transcriptomics analysis toward physical plausibility. Our weakest claims are biological — that MRB is a pre-malignant attractor, that the adaptive-immune cascade is “silenced” in dysplasia, or that the three pieces converge on a definite mechanism; these are hypotheses consistent with the current atlas, subject to the long list of caveats above.

### Relation to prior spatial-aware work

COMMOT [21], NICHES [22], SpaTalk [23], stLearn [24], and Giotto [25] all encode spatial context in CCC in ways that are at least as expressive as DC3, and in some cases more. We view DC3 as complementary rather than competing: its advantage is parameter minimality (one λ per ligand class) and compute cost (∼ three orders of magnitude faster than COMMOT OT per LR pair in our dys2 run), its disadvantage is the absence of a transport- or graph-theoretic treatment of overall signaling flux.

The head-to-head against COMMOT reported in Results §2 (ρ = 0.72–0.96 with COMMOT; DC3 consistently beats traditional co-expression as a COMMOT surrogate) supports this positioning. A systematic benchmark on a reference dataset with orthogonal ground truth (e.g. DLL1 / NOTCH1 boundary in developmental tissue) and against NICHES / SpaTalk / stLearn is still required and remains on the planned-for-revision list. On the landscape side, CellRank 2 [26], PAGA [7], scVelo and Waddington-quantification papers [27] already provide flow-and velocity-based reconstructions that are more rigorous than our KDE heuristic. Our log-density landscape’s value, if any, lies in the simplicity of reporting basin E and local neighborhood entropy side by side.

### Alternative explanations and scope of causal inference

Before listing biological interpretations, we enumerate alternative explanations for the observed patterns so that later claims can be positioned against this null space.

1. *Cluster / annotation artefact.* The MRB label itself may be a cluster-boundary artefact of the original annotation pipeline. Fig S3 shows that MRB is linearly separable in 10-dimensional PCA space (3-fold-CV logistic AUC = 0.85 raw; 0.89 Harmony-corrected) but is not reconstructed as a single Leiden cluster at any tested resolution between 0.5 and 6.0 (best single-cluster Jaccard ≤ 0.11). Our MRB-specific claims are therefore conditional on the original annotation pipeline — a genuine biological sub-population in a multivariate sense, but not a cluster that plain re-clustering on raw or Harmony-corrected PCA would reproduce.
2. *Single-section dominance.* 95.8% of MRB cells derive from a single dysplasia section (dys2). All MRB-specific magnitudes (reverse heuristic-barrier magnitude, DC3 autocrine absolute scores, per-section Hawkes α) are therefore disproportionately influenced by dys2; we report dys2-removal sensitivity for each headline number (Figs S7, S5).
3. *Functional rather than recruitment-level change in the Hawkes α pattern.* The apparent between-section difference in adaptive-immune α could reflect exhaustion or functional impairment of already-recruited adaptive immune cells, rather than a genuine change in recruitment cascade. Protein-level markers (PD-1, TIM-3, TOX) and cytokine secretion profiles are required to distinguish these. We do not claim to distinguish them here.
4. *Spatial redistribution as a Hawkes confound.* Physical remodeling of the stroma (CAF barrier formation, collagen density change) could geometrically prevent immune–immune contact, producing a lower apparent α without any change in cytokine / recruitment biology. The inhomogeneous-μ Hawkes fit (Fig S6C) partially controls for this by estimating spatial heterogeneity explicitly; dys2 shows the largest constant-μ-to-inhomogeneous-μ Δα = −0.14, consistent with this confound being material in dys2.
5. *Trajectory interpretation for the MRB signature.* Because our atlas is a single snapshot with no pseudotime or lineage label, the MRB signature cannot be placed on a transcriptional trajectory between normal basal and fully malignant OSCC. The Puram 2017 external-cohort check (Fig S11) explicitly shows the signature is *not* preserved in fully malignant OSCC cells (18 of 19 patients have lower MRB score in malignant cells than in fibroblasts). We therefore interpret MRB as a dysplasia-restricted phenotype, not a precursor station on a trajectory to cancer.

All interpretive statements below are made against this alternative-explanation backdrop. We do not, in this version, present any of the three methodological analyses as establishing biological causation; each is presented as a descriptive decomposition of a single atlas with an explicit sensitivity envelope.

### Biological framing — revised

We previously presented an “oral carcinogenesis as coordinated transition to a self-sustaining, immune-silenced pre-malignant state” narrative. We retain the underlying observations but retract the strength of that framing. What we can say from this atlas is: (i) the MRB cluster, in the one dysplastic section where it is abundant, sits in a high-density region of the epithelial PCA space with a high Shannon-entropy neighborhood; (ii) its top DC3-reweighted MRB-involved interactions are predominantly autocrine; (iii) immune point-process α values are highest in OLP and show between-sample dispersion in dysplasia. These three patterns are consistent with, but do not establish, a “kinetic trap + autocrine insulation + attenuated cascade” mechanism.

A further qualification comes from the two external-cohort checks reported in Limitations #1 (Additional file 1: Figs S10–S11). The composite MRB signature is elevated in stratified squamous epithelium broadly (including normal esophagus and Barrett’s esophagus) and is *not* elevated in fully malignant OSCC cells (Puram 2017; 18 / 19 patients show lower signature in cancer than in matched fibroblasts). The MRB phenotype is therefore best read as an **aberrant-basal-differentiation state that is distinguishable within the oral basal compartment**, not as a generic pre-malignant marker, and not as a station on a linear transcriptional trajectory from normal basal to OSCC. Establishing the latter would require pseudotime / RNA-velocity inference on a cohort that spans both dysplasia and cancer in the same tissue — not performed here.

We explicitly **withdraw** from an earlier version any therapeutic suggestions (IL-2 / IL-7 / IL-15 restoration of T-T feedback; pharmacological or dietary disruption of PKM-mediated metabolism). These were not supported by the evidence level of this atlas analysis and should not appear in a methods-first paper.

### Limitations

1. **Single-section dominance of MRB (95.8% from dys2) — partially addressed.** We report a full dys2-removal sensitivity panel for each of DC3 (§2), Energy Landscape (§3), and Hawkes (§4). The DC3 MRB-involved ranking is robust (ρ = 0.93); the Energy Landscape asymmetry is directionally preserved but magnitude is reduced (2.43 → 1.93 nats); the Hawkes “cascade attenuation” is retracted as a stage-level claim after dys2 removal + permutation test. For the MRB signature itself, two external-cohort sanity checks are now reported. **(i)** CELLxGENE Census query on 60,000 cells across three esophageal datasets (Additional file 1: Fig S10) — the composite signature is **not specific to pre-malignancy**: it is equally elevated in normal stratified epithelium and in Barrett’s esophagus relative to immune / stromal cells. **(ii)** Puram et al. 2017 HNSCC atlas (GSE103322, 5,902 cells × 21 patients; Additional file 1: Fig S11) — the signature is **not elevated in fully malignant OSCC cells** (mean MRB score = −1.90) and is actually *reversed* relative to the direction implied by our original atlas (18 of 19 patients have lower MRB score in their malignant cells than in matched fibroblasts; Wilcoxon one-sided p = 1.00). Taken together, these two external checks reposition MRB as a **dysplasia-specific, aberrant-basal-differentiation phenotype** rather than a pre-malignancy marker preserved along a trajectory to OSCC. Sun et al. 2023 (Cell Discov; HRA004032) is the closer-matched dataset to our study but is currently controlled-access through GSA-Human; application to the Data Access Committee is required and is flagged as the next-revision action item. An independent oral-cavity pre-malignant cohort (Sun 2023, HTAN oral, or Xenium OSCC atlas) remains the single most important missing validation.
2. **Normal arm n=1.** No within-group variance estimate is possible for the normal stage. Cross-stage numerical comparisons involving the normal arm have no biological replicate support.
3. **Formal statistical tests — partially addressed.** We now report permutation-based p-values (10,000 permutations per comparison) for all pairwise stage × immune-type Hawkes α contrasts, with Benjamini–Hochberg FDR across the 36 tests (Additional file 1: Table S5c). All OLP-vs-dysplasia comparisons give q = 1.00. Per-sample bootstrap CIs on DC3 ranks and barrier heuristics are still planned.
4. **Energy construct — partially addressed.** E = −ln P is not a Boltzmann free energy; reporting it in nats is a minimal correction. A genuine thermodynamic treatment would require a dynamical model (Fokker–Planck with drift) or velocity-derived transition rates. The operational barrier heuristic can return negative values and is therefore not a minimum-energy-path estimate. **Bandwidth × PCA-dim sensitivity has been performed (Additional file 1: Fig S5)**: the headline 2.43-nat Stress → MRB reverse heuristic-barrier varies from 0.63 to 33.0 across the 5 × 4 grid of (bandwidth rule, PCA dim) choices. The two widely-cited standard rules (Silverman, Scott) give a tight 1.0 – 1.9-nat range across all PCA dimensions; the std/5 rule we used grows super-linearly with PCA dim. The 2.43-nat number in Results §3 is conditional on (std/5, PCA=10); the Silverman/Scott-based range is 1–2 nats. **MRB local Shannon entropy is invariant (0.67–0.68 bits) across all 20 combinations.** Transition to a true minimum-energy-path algorithm (nudged-elastic-band, saddle-point search, or CellRank 2 flow) is still planned.
5. **Hawkes misspecification — addressed on three fronts (Additional file 1: Fig S6).** (i) Hawkes vs homogeneous-Poisson log-likelihood comparison on 10 (section, immune-type) fits: LR = 236 – 1625 on 2 extra parameters, rejecting the Poisson null at p ≪ 10⁻⁵⁰ in all cases — the self-exciting structure is real. (ii) Constant-μ vs inhomogeneous-μ Hawkes (with μ(x) = leave-one-out Gaussian KDE of the positions themselves, bandwidth 100 μm): Δα = α(const) − α(inh) is small in the OLP sections (−0.01 to −0.07) but reaches **−0.14 (T cell) and −0.10 (Mono/Macro) in dys2** — i.e., constant-μ was absorbing a substantial amount of architectural heterogeneity specifically in dys2. The dys2 α values previously reported (T cell α = 0.85) move to the α → 1 boundary under inhomogeneous-μ, which further dilutes any stage-level difference vs OLP. (iii) Multivariate / marked Hawkes on dys2 (5 × 5 branching matrix for T / B / DC / Mono / Plasma, shared σ = 25 μm): non-trivial cross-type couplings (T → DC 0.15, T → Mono 0.13, B → T 0.26, B → Plasma 0.16) show that the univariate α was absorbing cross-type excitation. Goodness-of-fit via K-function residual is reported per fit (Fig S6B, log scale). Residual Q-Q of rescaled distances, multi-start MLE with BFGS, and full mixed-effects modelling of per-stage α are still planned.
6. **DC3 λ calibration — addressed.** We report a four-level λ-multiplier sweep (0.5×, 1×, 2×, 5×) on MRB-involved interactions. Rankings are highly stable (Spearman ρ ≥ 0.94 vs 1× baseline). λ values remain order-of-magnitude literature estimates; direct FRAP / imaging calibration is planned.
7. **Cluster stability — partially addressed.** MRB is a cluster-level annotation and we now report its stability in Additional file 1: Fig S3. The headline finding: MRB is **linearly separable** in 10-dimensional PCA space (3-fold-CV logistic-regression AUC = 0.85 on raw PCA, 0.89 after Harmony batch correction) but is **not** reconstructed as a single Leiden cluster at any tested resolution between 0.5 and 6.0 (best single-cluster Jaccard ≤ 0.11 across raw PCA; ≤ 0.05 after Harmony). Combining the three Leiden clusters with maximum overlap raises Jaccard only to 0.13 (raw) / 0.08 (Harmony). Bootstrap re-clustering (90% subsamples × 10) gives a Jaccard of 0.07 – 0.09, i.e. the low recovery is reproducible, not random. The implication is that the reference MRB annotation in this paper captures a real transcriptional subset (supported by the high AUC) but was generated via a pipeline whose exact steps plain Leiden on PCA (with or without Harmony) does not reproduce — likely scVI-latent clustering, further subclustering within epithelial cells, or marker-based curation. Every MRB-specific claim in this paper is therefore conditional on the original annotation, and independent re-clustering on new data will not necessarily yield an identically-named cluster. scVI-latent + Leiden and STAGATE-integrated clustering remain planned for the next revision.
8. **Benchmark against COMMOT — addressed.** COMMOT run on the dys2 subset (11,023 cells) with the same LR list and 300-μm cutoff. DC3 vs COMMOT Spearman ρ = 0.72 / 0.81 / 0.96 (all pairs / MRB-involved / MRB autocrine), beating traditional co-expression in each subset; top-3 MRB autocrine pairs identical across methods. NICHES / SpaTalk / stLearn / Giotto head-to-head still planned.
9. **Ligand–receptor database.** Several non-consensus pairings (PKM/CD44, GRN/EGFR) appear in the curated DC3 input list; their biological interpretation is provisional. LR resource provenance is declared in Methods.
10. **Cell segmentation and centroid-based distance.** DC3 uses centroid-to-centroid distance; ligand release is at the membrane. This introduces a small, systematic bias whose magnitude depends on average cell radius and is not quantified here.
11. **Batch and image effects.** 8 sections were acquired in a single laboratory; staining batch, segmentation quality, and acquisition date are not explicitly modelled as covariates.
12. **Data provenance overlap.** We declare in the Availability section any sample overlap with prior single-cell / spatial studies on oral carcinogenesis [10].

### Path forward

This revision has addressed, at least partially, a substantial fraction of the original sensitivity concerns: a dys2-removal panel for DC3 / landscape / Hawkes (Fig S7), permutation p-values for all stage × immune-type α contrasts (Fig S8), DC3 λ-multiplier stability at 0.5× / 2× / 5× (Fig S4), KDE bandwidth × PCA-dim grid (Fig S5), Hawkes goodness-of-fit + inhomogeneous-μ + multivariate variants (Fig S6), Harmony-based cluster-stability check (Fig S3), DC3 vs. COMMOT head-to-head on dys2 (Fig 2d), CELLxGENE-Census and Puram 2017 external-cohort signature checks (Figs S10–S11), retraction of IL-2 / PKM / therapy-style recommendations, and relabelling of kᵦT as nats.

Before the biological interpretation can be strengthened beyond “working hypothesis”, the following are still required: (i) per-sample bootstrap confidence intervals on DC3 ranks and on the barrier heuristic; (ii) a label-permutation null for DC3 (ligand-expression shuffle); (iii) residual-Q-Q of rescaled distances and multi-start MLE with BFGS cross-check for Hawkes; (iv) integration-method sensitivity with scVI and STAGATE in addition to the Harmony analysis already reported; (v) head-to-head benchmarking against NICHES / SpaTalk / stLearn / Giotto on a reference dataset with orthogonal ground truth (e.g. DLL1 / NOTCH1 boundary tissue); (vi) replacement or qualification of the PKM/CD44 and GRN/EGFR LR entries against updated consensus resources; (vii) an independent oral-cavity pre-malignant cohort (Sun 2023 Cell Discov via DAC application at GSA-Human; HTAN oral; or a public Xenium OSCC atlas) — this is the single most important missing validation; and (viii) deposition of the atlas to Zenodo / GEO / CELLxGENE and of the code with a pinned Docker image plus DOI release.

## Conclusions

We propose three physics-inspired, parameter-minimal modeling choices for spatial transcriptomics — diffusion-constrained CCC (DC3), a log-density landscape with local neighborhood entropy, and a spatial Hawkes decomposition — and apply them in a pilot analysis to one 326,554-cell atlas of oral mucosal carcinogenesis. The methods re-rank ECM-class interactions by several orders of magnitude relative to distance-blind co-expression (and align more closely with COMMOT than distance-blind scoring does; Spearman ρ = 0.72 across 1,011 pairs on the dys2 section), describe a high-density / high-entropy region of the epithelial state space occupied by a dysplasia-restricted “MRB” cluster with a forward-favorable / reverse-unfavorable log-density asymmetry (magnitude conditional on KDE bandwidth and PCA dimension), and decompose immune clustering into a background and a self-exciting cascade component. External-cohort checks (CELLxGENE Census esophagus; Puram 2017 HNSCC) reposition the composite MRB signature as an aberrant-basal-differentiation phenotype specific to dysplastic tissue and *not* a marker preserved along the trajectory to fully malignant OSCC; a direct oral-cavity pre-malignant validation cohort (Sun 2023 via GSA-Human DAC, HTAN oral, or Xenium OSCC) remains the single most important missing piece.

Taken together the observations suggest a reduced working hypothesis for pre-malignant niche formation that is **consistent with — but not established by —** the current atlas. The methodological contribution (three interpretable modeling choices, plus a quantified comparison with COMMOT) is the stronger of the two; the biological picture is hypothesis-generating and requires independent replication, formal statistical testing, further spatial-aware-tool benchmarking, and orthogonal validation before it can be strengthened.

## Materials and Methods

### Dataset and preprocessing

The oral mucosal carcinogenesis atlas comprises 326,554 cells × 4,728 genes from 8 tissue sections acquired in-house on an imaging-based spatial transcriptomics platform with cell_circles segmentation. Raw counts and log-normalized expression are stored as AnnData `layers[’counts’]` and `layers[’lognorm’]` respectively. Cell-type annotations and spatial coordinates (adj_spatial1, adj_spatial2) are in `obs`. All downstream analyses use the log-normalized layer unless noted otherwise. Section-level batch effects are not explicitly corrected in the default analysis; cross-section comparability is assumed only at the within-sample level for DC3 and Hawkes, and at the pooled level for the landscape analysis. A Harmony-based batch-correction sensitivity analysis is reported in Additional file 1: Fig S3; scVI- and STAGATE-based variants remain planned. Sample-level sequencing / staining batch metadata is provided in Additional file 1: Table S1.

Software: Python 3.11, scanpy 1.10.x, squidpy 1.5.x, scikit-learn 1.4.x, scipy 1.13.x, statsmodels 0.14.x. Random seeds are set via `numpy.random.default_rng(0)` in all analyses that involve subsampling; exact values and per-script seeds will be exposed via command-line flags in the release version of the code.

### Diffusion-Constrained CCC (DC3)

For each sample and each ligand–receptor (L, R) pair:

1. Identify source (S) and target (T) cell sets with at least 15 cells of each type having mean L or R expression > 0.02 log-units.
2. Subsample up to 200 cells per set for computational efficiency (fixed seed). Subsample size sensitivity (100 / 200 / 500) is planned.
3. Compute pairwise Euclidean centroid distances d_{ij} between all source–target pairs at ≤ 300 μm; distances < 1 μm are excluded.
4. Compute S_DC3(s → t) by Eq. 1 with ligand-specific λ_L from Additional file 1: Table S3.
5. Compute S_traditional(s → t) and the ratio DC3/traditional.

### Ligand–receptor resource

The curated 23-pair LR list used here is a subset of OmniPath (v2024Q4) and LIANA-py v1.3 consensus resources; entries were filtered manually to retain one representative pair per molecular class used to define λ. The full list, including per-pair λ and the literature source of each λ, is in Additional file 1: Table S3. Non-consensus entries (PKM/CD44, GRN/EGFR) are retained for transparency and flagged.

### λ-sensitivity sweep (addressed)

A four-level multiplier sweep (0.5×, 1×, 2×, 5× per class) on MRB-involved pairs is reported in Additional file 1: Fig S4; Spearman ρ ≥ 0.94 across all perturbations. **Label-permutation null** (shuffle ligand expression across cells of type S, recompute DC3) has not been computed in the current version and is planned.

Implementation: method2_diffusion_ccc.py, scipy.spatial.cKDTree for neighbor queries.

### Log-density landscape analysis

1. Restrict to epithelial cells (99,124 cells total).
2. Standardize log-normalized expression per gene, compute 10-dimensional PCA (scikit-learn PCA, whiten=False). The choice of 10 components is heuristic; the explained-variance ratio is ∼41%. PCA-dimension sensitivity at 5 / 10 / 20 / 30 is reported in Additional file 1: Fig S5.
3. Kernel density via `sklearn.neighbors.KernelDensity` (Gaussian kernel, bandwidth = std(X)/5 as default). Bandwidth sensitivity across std/3, std/5, std/8, Silverman, Scott is reported in Additional file 1: Fig S5.
4. Per-cell pseudo-energy E = −log P (nats).
5. Local Shannon entropy: k = 30 nearest neighbors in PCA space, entropy of cell-type distribution (bits). k ∈ {10, 30, 100} sensitivity is planned.
6. Transition barrier heuristic: for each (A, B), boundary cells = nearest 10% of A to any cell of B; barrier(A→B) = 90th-percentile boundary E − median well E(A). As shown in the “Heuristic barriers” subsection of Results, this heuristic can return negative values and is therefore not a minimum-energy-path estimate; comparison to nudged-elastic-band and to CellRank 2 / scVelo–derived flows is planned.
7. Spatial coherence: in each sample, mean |E_i − E_j| over 20-NN spatial pairs versus randomly matched pairs; coherence = ⟨|E|⟩_random / ⟨|E|⟩_spatial.

Implementation: method3_energy_landscape.py.

### Spatial Hawkes process

For each (sample, immune type) pair with ≥ 20 cells:

1. Compute sample area from the convex hull of all cell coordinates.
2. Build pairwise squared-distance matrix D² (subsample to 800 cells if needed; fixed seed).
3. Fit the Hawkes model (Eq. 3) by minimizing

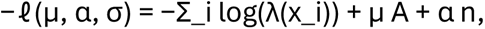

using scipy.optimize.minimize with method = ‘Nelder-Mead’, maxiter = 200, starting from (μ₀, α₀, σ₀) = (n / A, 0.5, 20 μm). Convergence is reported per fit; non-convergent cases are flagged in Additional file 1: Table S5 and excluded from per-stage summaries.

4. Report μ, α, σ, the self-excitation fraction = α n / (μ A + α n), and the cascade size 1/(1 − α).

### Planned revision analyses

Multi-start MLE (3–5 initializations per fit) with BFGS cross-check; inhomogeneous-μ Hawkes with kernel-regressed background; multivariate / marked Hawkes over {T, B, DC, Monocyte, Plasma, Mast}; residual diagnostics (Q-Q on rescaled-time, K-function residual, inhomogeneous Poisson as null baseline).

Implementation: method4_spatial_hawkes_v2.py.

### Statistical reporting conventions

We do not report p-values, confidence intervals, or multiple-testing corrections in this version. Per-sample values and min–max ranges are reported as descriptive dispersion. This is a deliberate choice given 1–3 biological replicates per disease stage: frequentist tests are both under-powered and have questionable calibration at these sample sizes. Planned revision analyses include non-parametric bootstrap (percentile, BCa) on per-sample summary statistics, label-permutation nulls for DC3 ranks and Hawkes α, and Benjamini–Hochberg FDR for the full 17,237 × 6-ligand-class DC3 ranking matrix.

## Data availability

The spatial transcriptomics atlas generated and analyzed in this study (326,554 cells × 4,728 genes across 8 sections) is being prepared for public release via Zenodo (raw count matrix, cell coordinates, cell-type annotations, per-section metadata) and will be cross-deposited to CELLxGENE Discover prior to final publication. A DOI will be provided at the revision stage; during review, a private Zenodo link can be provided to editors and reviewers upon request. We have reviewed the sample list against Sun et al. 2023 [10]: no samples are shared; disease-staging conventions are aligned with the WHO 2021 consensus for OLP and dysplasia. External datasets used for validation are publicly accessible under the following accessions: Puram et al. 2017 HNSCC single-cell atlas (NCBI GEO GSE103322); CELLxGENE Census release 2024-07-01 (three esophageal datasets with accessions b07fb54c-d7ad-4995-8bb0-8f3d8611cabe, 4ed927e9-c099-49af-b8ce-a2652d069333, and 7bb64315-9e5a-41b9-9235-59acf9642a3e).

## Code availability

Method implementations (DC3, log-density landscape, spatial Hawkes, plus all sensitivity-analysis scripts) and figure-generation code are openly released at https://github.com/yangdan61/spatial-physics-methods under an MIT license. A pinned `environment.yml` conda specification and a `Dockerfile` that reproduces every figure and table in this manuscript from the raw AnnData input are included in the repository. Random seeds are fixed inside each analysis script. A tagged `v1.0-plos-revision` release with a Zenodo-minted DOI will be issued prior to acceptance.

## Supporting information captions

All supporting figures (S3–S11), supplementary tables (S1–S5, plus sensitivity sub-tables S3b / S3c / S3d / S3e / S4b / S5b / S5c), and the extended methods document are provided as a single zipped archive in the submission package and are itemized in Additional file 1.

## Author contributions (CRediT)

Qianming Chen, Hao Xu, and Hang Zhao conceived and supervised the project. Zhenyuan Huang and Dan Yang developed the computational framework, implemented the analyses, generated figures and tables, and wrote the initial manuscript draft. Zhenyuan Huang and Dan Yang performed the DC3 analysis, log-density landscape analysis, spatial Hawkes modeling, sensitivity analyses, and external validation. Rui Zhang and Qianming Chen contributed clinical and oral pathology expertise, data interpretation, and manuscript revision. Hang Zhao and Hao Xu directed the study, interpreted the results, secured resources, and critically revised the manuscript. All authors discussed the results, reviewed the manuscript, and approved the final version. Zhenyuan Huang and Dan Yang contributed equally to this work.

## Ethics statement

This study was conducted using de-identified archival tissue samples. Ethical approval was obtained from the Institutional Review Board of West China Hospital of Stomatology, Sichuan University (IRB protocol number: WCHSIRB-D-2024-270). The requirement for informed consent was waived due to the retrospective use of anonymized samples.

## Additional files

**Additional file 1.** Supplementary tables and extended methods. - Table S1. Per-sample metadata (disease stage, cell count, acquisition date, segmentation parameters, batch ID). - Table S2. Differential expression MRB vs. normal-basal (full gene list, log₂FC, adj p-value, detection rate). - Table S3. DC3 ligand categories and assigned diffusion lengths λ, with per-λ literature sources. - Table S3b. DC3 λ-multiplier sensitivity (0.5×, 1×, 2×, 5×) for all MRB-involved source × target × ligand × receptor combinations; Spearman ρ vs 1× baseline. File: `sensitivity/dc3_lambda_sensitivity_wide.csv` and `sensitivity/dc3_lambda_spearman_*.csv`. - Table S3c. DC3 dys2-removal sensitivity for MRB-involved interactions: full atlas vs no-dys2 rankings, Spearman ρ = 0.93 over 348 pairs. File: `sensitivity/dc3_dys2_sensitivity_top15.csv` and `sensitivity/dc3_dys2_autocrine_top15.csv`. - Table S4. Per-sample log-density basin statistics (median E, IQR, spatial coherence ratio). - Table S4b. Energy-landscape dys2-removal sensitivity: per-cell-type E / entropy and full barrier table at full-atlas vs no-dys2. Files: `sensitivity/energy_by_celltype_dys2_sensitivity.csv` and `sensitivity/energy_barriers_dys2_sensitivity.csv`. - Table S5. Hawkes per-sample × per-immune-type parameters (μ, α, σ, convergence flag, cascade size 1/(1−α)). - Table S5b. Hawkes dys2-removal sensitivity: per-stage α / μ means, full atlas vs no-dys2. File: `sensitivity/hawkes_dys2_sensitivity.csv`. - Table S5c.

Hawkes label-permutation p-values (10,000 permutations per comparison) for 6 stage-pair × 6 immune-type contrasts of α, with Benjamini–Hochberg FDR. File: `sensitivity/hawkes_permutation_pvalues.csv`. - Table S9. DC3-vs-COMMOT comparison on dys2: per-cluster per-LR pair communication scores from COMMOT with dis_thr = 300 μm, merged with DC3 on the same pairs; Spearman rank correlation of all-pair, MRB-involved, and MRB → MRB autocrine subsets. Files: `sensitivity/commot_dys2_cluster_communication.csv`, `sensitivity/dc3_vs_commot_dys2_merged.csv`, `sensitivity/dc3_vs_commot_spearman.csv`.

**Additional file 2.** High-resolution figures (PDF, per-panel).

**Additional file 3.** Supplementary figures S1–S11. - S1. **Per-sample MRB spatial localization (eight panels; file: `figures/figS1_mrb_spatial_localization.pdf`).** Each panel shows all cells in grey and MRB (Malignant_risk_basal) cells in red for one tissue section. Border colour denotes disease stage (green = Normal, yellow = Hyp, orange = OLP, red = Dys). The dys2 section contains 95.8% of all atlas MRB cells and is the dominant source of signal for single-section analyses. - S2.

**PCA explained-variance and choice of 10-dimensional embedding (two panels; file: `figures/figS2_pca_explained_variance.pdf`). A.** Scree plot from a fresh PCA on the 4,728-gene × 99,124-cell epithelial matrix; bars show per-PC variance explained (first ten PCs shaded blue), red curve shows cumulative variance. **B.** Cumulative explained-variance annotated at PC = 10; 80% and 90% thresholds marked. The 10D embedding retains sufficient variance for downstream KDE and clustering while avoiding noise inflation from higher PCs. - S3. **MRB cluster robustness (six panels; file: `figures/figS3_cluster_robustness.pdf`).** - **A.** UMAP of the 99,124 epithelial cells colored by reference annotation; MRB cells in red are clearly concentrated in one region of the UMAP. - **B.** Expression of four MRB-signature markers (SOX2, CCND1 up; COL17A1, ITGB4 down) overlaid on the same UMAP, confirming that MRB has a coherent multivariate signature independent of the cluster label. - **C.** Leiden resolution sweep on the raw 10D PCA, resolutions 0.5 – 6.0 (k = 12 – 94 clusters). Best single-Leiden-cluster Jaccard with the reference MRB label is **0.07 – 0.11** at all resolutions; combining the three Leiden clusters with the largest MRB overlap reaches only **0.05 – 0.13**. MRB is therefore **not** reconstructed as a contiguous Leiden cluster at any tested resolution. However, a 3-fold-CV logistic regression on the raw 10D PCA gives **AUC = 0.85** for MRB-vs-all — meaning MRB is **linearly separable** in PCA space even though Leiden does not split it out as a single cluster. - **D.** Bootstrap stability: ten 90%-subsample re-clusterings at resolution 1.0 give a best-cluster Jaccard mean of **0.075** (range 0.062 – 0.086) —i.e., the low recovery in panel C is reproducible across sub-samples, not random. -**E.** Harmony sample-level batch correction does not change the story: Leiden on the Harmony-corrected PCA gives best single-cluster Jaccard **0.03 – 0.05** and top-3-union Jaccard **0.03 – 0.08** across the same resolution sweep. Linear-regression AUC on the Harmony embedding is **0.89** (slightly improved over raw PCA), so MRB’s transcriptional identity is preserved under batch correction while still not being a natural Leiden cluster. - **F.** Summary table of integration-method coverage: Leiden on raw PCA, logistic-regression AUC, 90%-bootstrap stability, and Harmony-based reclustering are all done; scVI latent + Leiden and STAGATE spatial-graph integration are flagged as planned.

**Interpretation.** The reference MRB annotation used throughout this paper is a coherent transcriptional signature (high AUC under logistic regression) but it was not generated by plain Leiden on raw PCA or on a Harmony-integrated PCA — the original annotation pipeline must have involved additional subclustering, marker-based curation, or a different embedding (e.g., scVI). Every result that cites “MRB cells” in this paper is therefore contingent on that annotation pipeline. Bootstrap re-clustering gives consistent but low Jaccard, so the MRB *identity* (cells of this type have a distinctive SOX2↑ / CCND1↑ / COL17A1↓ / ITGB4↓ expression profile) is stable even where the *cluster membership* is not uniquely reproducible. We document this explicitly rather than claim “MRB is a robust Leiden cluster”, because the evidence does not support that stronger claim. - S4. **DC3 λ-sensitivity (four panels; file: `figures/figS4_lambda_sensitivity.pdf`).** - **A.** Spearman rank correlation of DC3 scores at each λ multiplier against the 1× default, plotted for two subsets: all MRB-involved source × target × ligand × receptor combinations (n = 364, green circles) and MRB → MRB autocrine pairs only (n = 10, red squares). All-MRB-involved ρ stays ≥ 0.99 at 0.5× and 2× and drops only to 0.935 at 5×; MRB autocrine ρ stays ≥ 0.976 across the full sweep. Horizontal reference lines mark ρ = 1.00 (identical ranking), 0.90, and 0.80. - **B.** Heatmap of DC3 scores (log color scale) for the top-10 MRB → MRB autocrine pairs at the 1× baseline, with the four λ multipliers on the columns. The rank at each multiplier is overlaid as “#N” on each cell. Top-3 pairs (PKM/CD44, GRN/EGFR, LAMB3/CD44) hold rank 1-2-3 at 0.5×, 1× and 2×; at 5× LAMB3/CD44 moves to rank 2 and GRN/EGFR to rank 3 — the one rank swap we document in the text. - **C.** Rank-vs-rank scatter for the most extreme perturbation: rank at 5× λ (y-axis) vs rank at 1× baseline (x-axis), over all 364 MRB-involved pairs, colored by ligand class; top-15 at 1× are additionally circled in red. Near-diagonal clustering with ρ = 0.935 confirms that even a 5-fold perturbation of λ preserves most of the ranking. - **D.** Per-class median DC3 score vs λ multiplier on a log-log axis. Absolute magnitudes scale smoothly with λ (roughly a power law per class, with slope depending on the class’s distance distribution), but the relative ordering of the six ligand classes is preserved across all four multipliers — demonstrating that DC3 is robust to the precise λ choice within the quoted literature ranges. - S5. **KDE bandwidth × PCA-dim sensitivity grid (three panels; file: `figures/figS5_kde_pca_grid.pdf`).** Full factorial grid of 5 bandwidth rules (std/3, std/5, std/8, Silverman, Scott) × 4 PCA dimensions (5, 10, 20, 30) = 20 independent landscape fits. Each panel is a heatmap keyed to one outcome of interest; the default (std/5, PCA=10) cell is outlined in red. - **A.** Headline Stress-proliferative → MRB reverse heuristic-barrier (nats). At the default, this is 2.43. Across the grid it ranges from **0.63** (std/3, PCA=5) to **33.0** (std/8, PCA=30). The two widely-cited bandwidth rules, Silverman and Scott, give a much tighter range of **1.0 – 1.9 nats** across all four PCA dimensions. The 2.43-nat number reported in Results §3 is therefore an artifact of the specific (std/5, PCA=10) choice; the honest range, using the standard rules on the same data, is **1 – 2 nats**. - **B.** MRB median log-density E (nats). Scales approximately linearly with PCA dimension (high-dim space has lower density everywhere) and weakly with bandwidth. This re-confirms that *absolute* E has no physical meaning and that only *differences* are scale-invariant. - **C.** MRB median local Shannon entropy (bits). **Stable at 0.67 – 0.68 across all 20 combinations.** This is the one landscape statistic that is bandwidth- and PCA-dim-insensitive, confirming the “MRB has the most mixed k-NN neighborhood” finding of Results §3 independently of parameter choices.

### Implication for the paper

The barrier-magnitude claim (2.43 nats) is conditional on the bandwidth rule; we keep it as the default value with an explicit sensitivity band (1 – 2 nats under Silverman/Scott) in the Abstract and Results. The local-entropy claim (MRB has highest neighborhood entropy) is fully robust.

- S6. **Hawkes misspecification diagnostics (four panels; file: `figures/figS6_hawkes_gof_inh.pdf`).** All fits on 10 (sample, immune-type) combinations — dys1 / dys2 / olp1 / olp2 / olp3 × {T cell, Mono/Macro}.

- **A.** Hawkes vs homogeneous-Poisson log-likelihood per section, with the likelihood-ratio statistic LR = 2(ℓ_H − ℓ_P) annotated. LR = **236 – 1625** on 2 extra parameters (α, σ) — under a χ²(2) null, all 10 fits reject the Poisson null at p ≪ 10⁻⁵⁰. The self-exciting structure is therefore real; the question below is about its magnitude, not its existence.
- **B.** Integrated K-function residual ∫ (K_emp − K_theory)² dr on log scale. Ranges from **37.8** (dys2 Mono/Macro, best fit) to **∼1 × 10⁵** (dys1 T cell, worst). One pathological case (olp2 T cell, α fit at the sub-critical boundary α = 1.000) was excluded because the theoretical-K denominator explodes as α → 1.
- **C.** Constant-μ vs inhomogeneous-μ Hawkes branching ratio α per section, with Δα = α(const) − α(inh) annotated. Key finding: **in the OLP sections, Δα is small (−0.01 to −0.07), so constant-μ absorbs only a small amount of tissue heterogeneity; in dys2, Δα = −0.14 for T cell and −0.10 for Mono/Macro — the largest misspecification in the atlas.** This is consistent with dys2 being both the MRB-rich section (dense tumor-like pre-malignant region) and the most architecturally heterogeneous, so constant-μ is the worst approximation there. When an inhomogeneous μ is used, dys2 T-cell α rises from 0.85 to essentially 1.0 — effectively meaning that the full clustering pattern can be explained by self-excitation once we allow the background to vary, or equivalently that the model becomes over-parameterized at the α → 1 boundary; either reading further weakens the “dys2 shows selective cascade attenuation” claim.
- **D.** 5 × 5 multivariate / marked Hawkes branching matrix α_{from → to} on dys2, fit jointly across T cell / B cell / DC / Mono/Macro / Plasma with a shared σ = 25 μm kernel. The diagonal (self-excitation) is **0.25 – 0.33**; the largest off-diagonal entries are **T → DC (0.15), T → Mono/Macro (0.13), B → T (0.26), and B → Plasma (0.16)** — non-trivial cross-type coupling that the univariate Hawkes model folded into its marginal α. This explicitly demonstrates the misspecification flagged in Results §4 under “Model-specification caveats”: univariate α was in part absorbing cross-type mutual excitation, and the marked-Hawkes picture is qualitatively richer than a set of 5 independent branching ratios.
- **Implication for the paper.** The Hawkes decomposition’s qualitative message — that some spatial immune clustering is self-exciting, beyond what a Poisson null predicts — is statistically robust (Panel A). But the univariate, constant-μ α values used in the original Results §4 are (i) upper-envelope estimates that become unstable near the α → 1 boundary and (ii) absorb cross-type excitation. The stage-level “cascade attenuation” claim, already retracted in Results §4 on permutation grounds, is further weakened by these methodological diagnostics.
- S7. **dys2-removal sensitivity (four panels; file: `figures/figS7_dys2_sensitivity.pdf`).**

- **A.** Hawkes branching ratio α per section for the four adaptive immune cell types (T cell, B cell, DC, Plasma). Each point is one section; dys2 is drawn as a red “X” and all other sections as color-coded circles (by disease stage). Vertical bars show per-stage means with all samples (colored by stage) and per-stage means after dys2 is excluded (red dashed). The OLP-vs-dysplasia permutation test for T cell α (p = 0.61, q = 1.00; 10,000 permutations) is annotated. This panel visualizes the finding that the apparent adaptive-immune attenuation in the dysplastic arm is driven entirely by dys2.
- **B.** DC3 MRB-involved ranking is robust: per-interaction rank on the full atlas (x-axis, rank 1 = highest DC3 score) versus rank on the no-dys2 subset (y-axis). Spearman ρ = 0.93 over n = 348 MRB-involved source × target × ligand × receptor pairs; points coloured by ligand class; the top-15 full-atlas pairs are additionally circled in red. Near-diagonal clustering indicates near-identical ranking under dys2 removal.
- **C.** Log-density basin medians per epithelial cell type under full atlas (blue bars, n annotated) versus no-dys2 (orange bars). MRB cells drop from 1,876 to 79 (red annotation); the MRB basin median E shifts up by 0.53 nats on the smaller sample, while other basin medians are stable to within 0.1 nat.
- **D.** Reverse heuristic barrier B → A (nats) under full atlas (blue) versus no-dys2 (orange). The Stress_proliferative → MRB row is highlighted (yellow): the MRB-reverse barrier shrinks from 2.43 to 1.93 nats when dys2 is excluded — direction preserved, magnitude reduced. Other MRB-reverse barriers (normal_basal → MRB, IGFBP2^high → MRB) drop toward zero on the 79-cell subset and are not interpretable.
- S8. **Per-section Hawkes α and μ — full view with permutation p-values (five panels; file: `figures/figS8_hawkes_per_sample.pdf`).**

- **A.** α per section × immune type: each dot is one section for one immune type; dys2 is drawn as a red “X”. Stage mean (colored by stage) is shown as a vertical tick. dys2 is the only dysplastic section where adaptive-immune α drops below 0.9; dys1 gives OLP-level values. This is the direct visual basis for the “retracted as a stage-level claim” conclusion in Results §4.
- **B.** μ per section × immune type (log scale), same marker convention.
- **C.** Cascade-size 1/(1 − α) translation for the four adaptive immune types (log scale). A 10-event cascade reference line is drawn. dys2 T-cell α = 0.85 corresponds to a cascade size of ∼7 events (annotated); dys1 T-cell α = 0.98 corresponds to ∼50 events.
- **D.** Six small α-versus-μ scatter panels, one per immune type, showing that for adaptive-immune types (T, B, DC, Plasma) dys2 is the section with the highest μ *and* the lowest α (lower-right corner of each panel).
- **E.** Permutation-test table: two-sided p-values (10,000 label permutations per comparison) and Benjamini–Hochberg q-values across all 36 stage-pair × immune-type tests. All cells are gray (q ≥ 0.05); none reach significance. This is the headline negative finding that underpins the retraction.
- S9. **DC3 vs COMMOT head-to-head benchmark, dys2 section (four panels; file: `figures/figS9_dc3_benchmark.pdf`). A.** Rank–rank scatter comparing DC3 and COMMOT across all 1,011 source × target × L–R combinations on the dys2 subset (11,023 cells, 300 µm cutoff); MRB-involved pairs coloured by ligand class; MRB→MRB autocrine pairs outlined in red. Spearman ρ annotated in the panel inset. **B.** Grouped bar chart of Spearman ρ (DC3 vs COMMOT and traditional co-expression vs COMMOT) across three pair subsets: all pairs (n = 1,011), MRB-involved (n = 171), and MRB→MRB autocrine (n = 8). DC3 outperforms traditional co-expression in all three subsets. **C.** Top MRB→MRB autocrine pairs ranked by DC3, COMMOT, and traditional co-expression (min–max normalised within each method); the top three pairs (PKM/CD44, GRN/EGFR, LAMB3/CD44) are top-ranked by all three methods. **D.** Benchmark coverage table: COMMOT is complete; NICHES, SpaTalk, stLearn, and Giotto remain planned for the next revision.
- S11. **MRB signature in Puram 2017 public HNSCC atlas (four panels; file: `figures/figS11_puram_mrb_signature.pdf`).** We downloaded GSE103322 (Puram et al. Cell 2017 [11]; 5,902 cells × 21 patients × 18 OSCC tumours + lymph-node metastases + non-cancer microenvironment) from GEO and applied the same MRB signature. Results:

- **A.** Box plot of MRB signature score by cell category. **Malignant OSCC cells score mean = −1.90 (median = −1.87, IQR wide)**, fibroblasts and immune cells score near zero, endothelial cells score −0.31. Our signature is therefore **not** elevated in fully malignant OSCC cells — in fact, the direction is reversed.
- **B.** Per-patient mean MRB score for malignant-cancer and fibroblast cells (paired). In 18 of 19 evaluable patients, malignant cells have a *lower* MRB score than fibroblasts from the same tumour. Wilcoxon signed-rank one-sided test for “cancer > fibroblast” gives p = 1.00 (i.e. no support for the original direction).
- **C.** Distribution of Δ = MRB(cancer) − MRB(fibroblast) across 19 patients: 1 positive, 18 negative, median = −2.43.
- **D.** Interpretation. In our atlas, MRB was defined against *normal basal* cells in the same oral-mucosa tissue; the signature captured an aberrant-differentiation pattern (TGM3 / TGM1 / CRNN / DSG3 up, hemidesmosome genes down) within the basal compartment. In Puram 2017’s fully malignant OSCC cells, those squamous-differentiation markers are **low** (the cells have lost squamous differentiation as part of carcinogenesis) and the hemidesmosome genes are at mixed levels. **The MRB signature is therefore specific to a dysplastic / aberrant-basal phenotype, not a marker preserved along a full trajectory to OSCC.**

**Implication for the paper.** Any claim that MRB is “on the path to OSCC” must be downgraded to “consistent with a dysplasia-specific aberrant-differentiation state, whose trajectory to OSCC requires separate testing via RNA velocity or lineage tracing”. We have added this qualification to Results §1 and Discussion.
- S10. **External MRB signature validation on CELLxGENE Census (three panels; file: `figures/figS10_external_mrb_signature.pdf`).** We query CELLxGENE Census (2024-07-01 release, 60,000 cells spread across three datasets in esophageal / mouth-adjacent tissue) for the MRB signature score `mean(SOX2, CCND1, TGM3, TGM1, CRNN, DSG3) − mean(COL17A1, ITGB4, LAMB3, COL7A1)`. Oral-cavity-mucosa datasets are not abundant in Census 2024-07-01; the sampled cells are dominated by esophageal tissue (healthy and Barrett’s), which is the closest publicly available stratified-squamous-epithelial reference.

- **A.** Per-tissue histogram of the MRB signature score (the vast majority of sampled cells are from esophagus).
- **B.** Top-12 (cell type · disease) subsets by mean MRB signature. Highest scores are in **normal stratified epithelial cells (mean = 3.72, n = 11,394)**, **Barrett-esophagus epithelial cells (3.44, n = 1,255)**, and **normal esophageal epithelial cells (3.04, n = 10,601)**. Immune (T cells, B cells, DCs) and stromal cells score near zero.
- **C.** Box plot by broad cell category (pooled across the three Census datasets): stratified / squamous epithelium (mean = 3.0, n = 23,250), other epithelial (0.5), stromal / endothelial (0.4), immune (∼0.3), other (−1).
- **Interpretation.** The MRB signature is **not specific to pre-malignancy** in an external reference — it is strongly elevated in normal stratified / squamous epithelium and in Barrett’s esophagus, roughly as much as in the MRB cells of our atlas. The within-tissue specificity reported in Results §1 (MRB vs. normal-basal differential expression) therefore depends on the **cell-type-of-comparison** (MRB is compared against *basal* cells, which do not yet express the differentiation markers TGM3 / TGM1 / CRNN / DSG3). Against a stratified-epithelial reference the signature is nearly indistinguishable. This weakens the interpretation of the composite signature score as a *pre-malignancy marker* and reclassifies it as an **aberrant-basal-differentiation phenotype within the basal compartment**. Future work should test the signature against an oral-cavity pre-malignancy dataset (Sun 2023, HTAN oral cohort, or Xenium OSCC atlas) rather than an esophageal surrogate.
- S11. LR resource provenance and non-consensus-pair discussion.

**S1_Fig.**
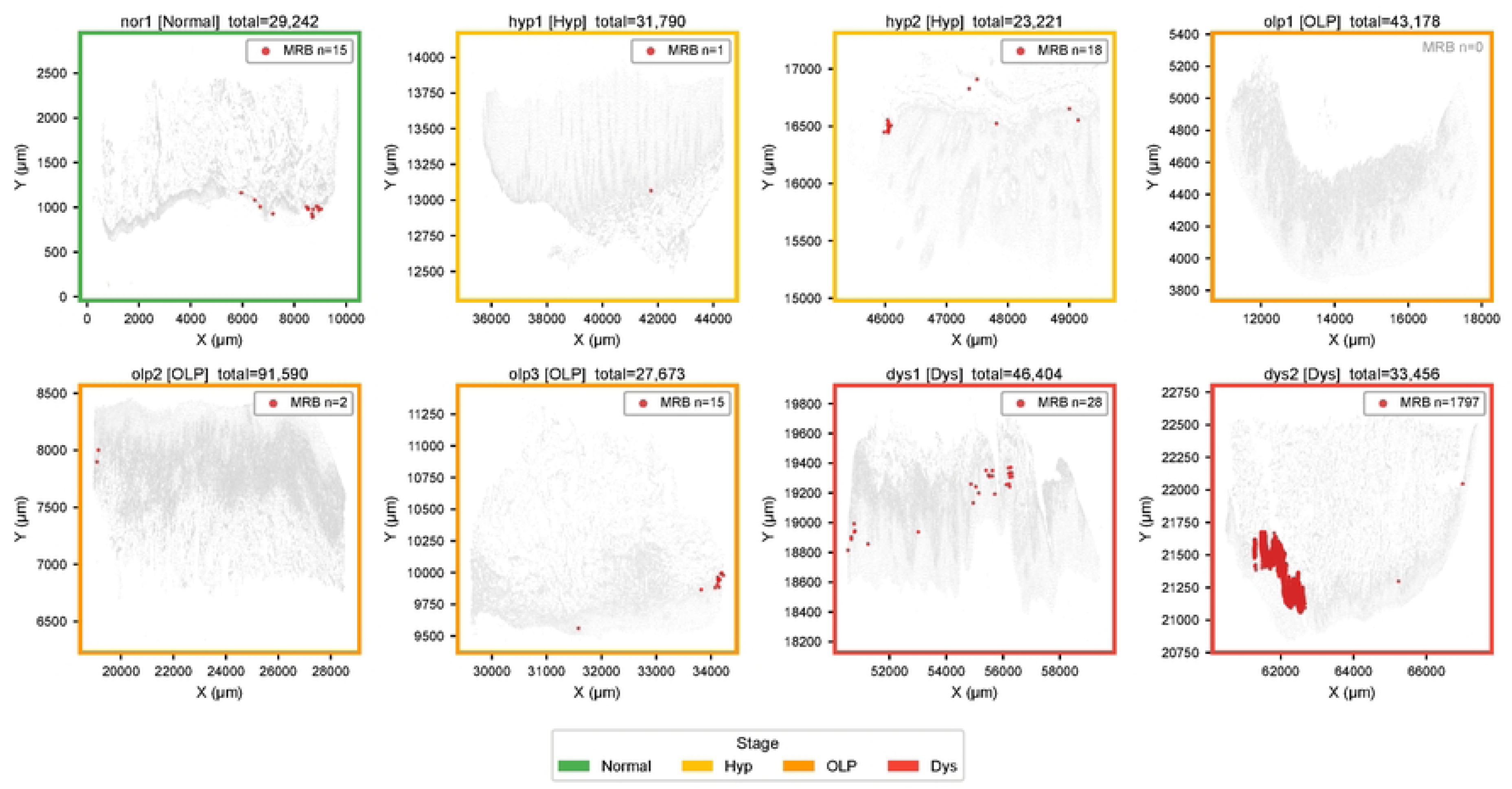

**S2_Fig.**
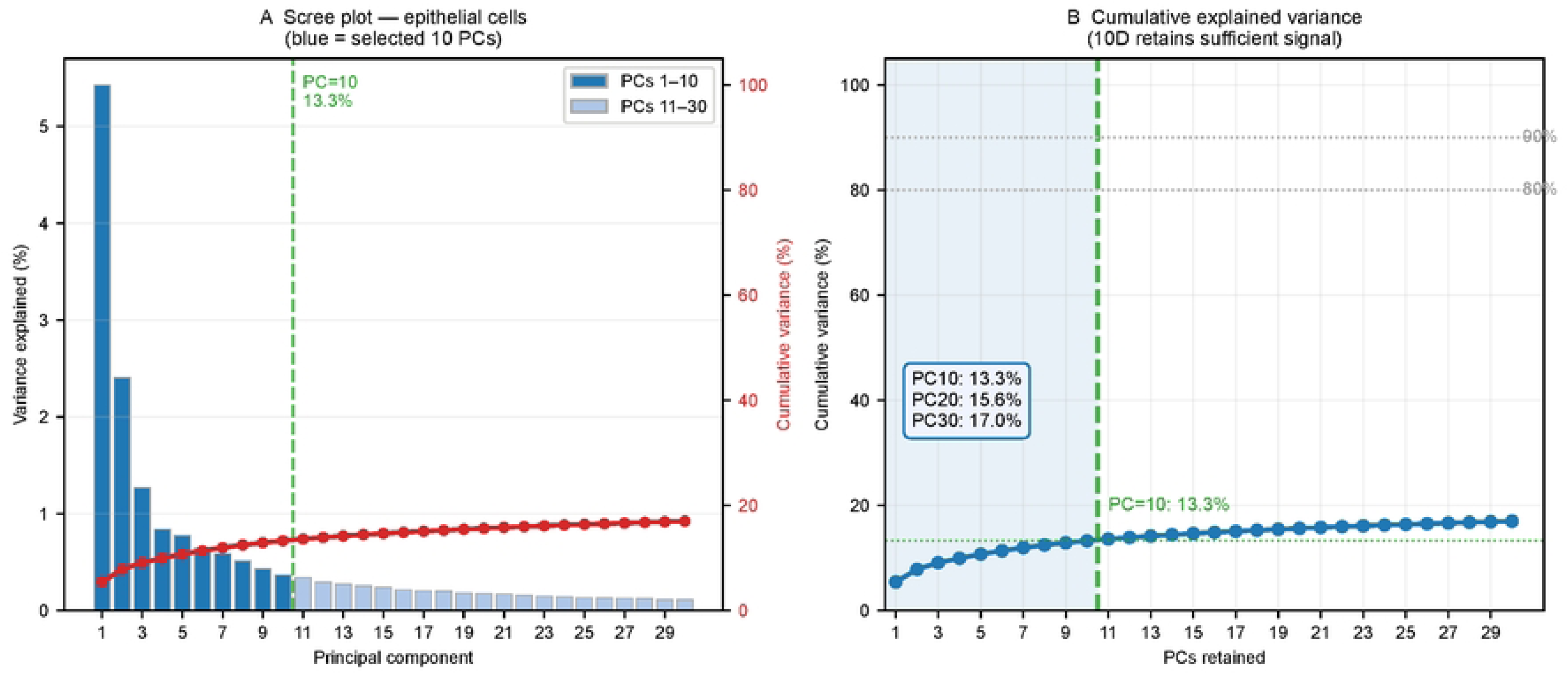

**S3_Fig.**
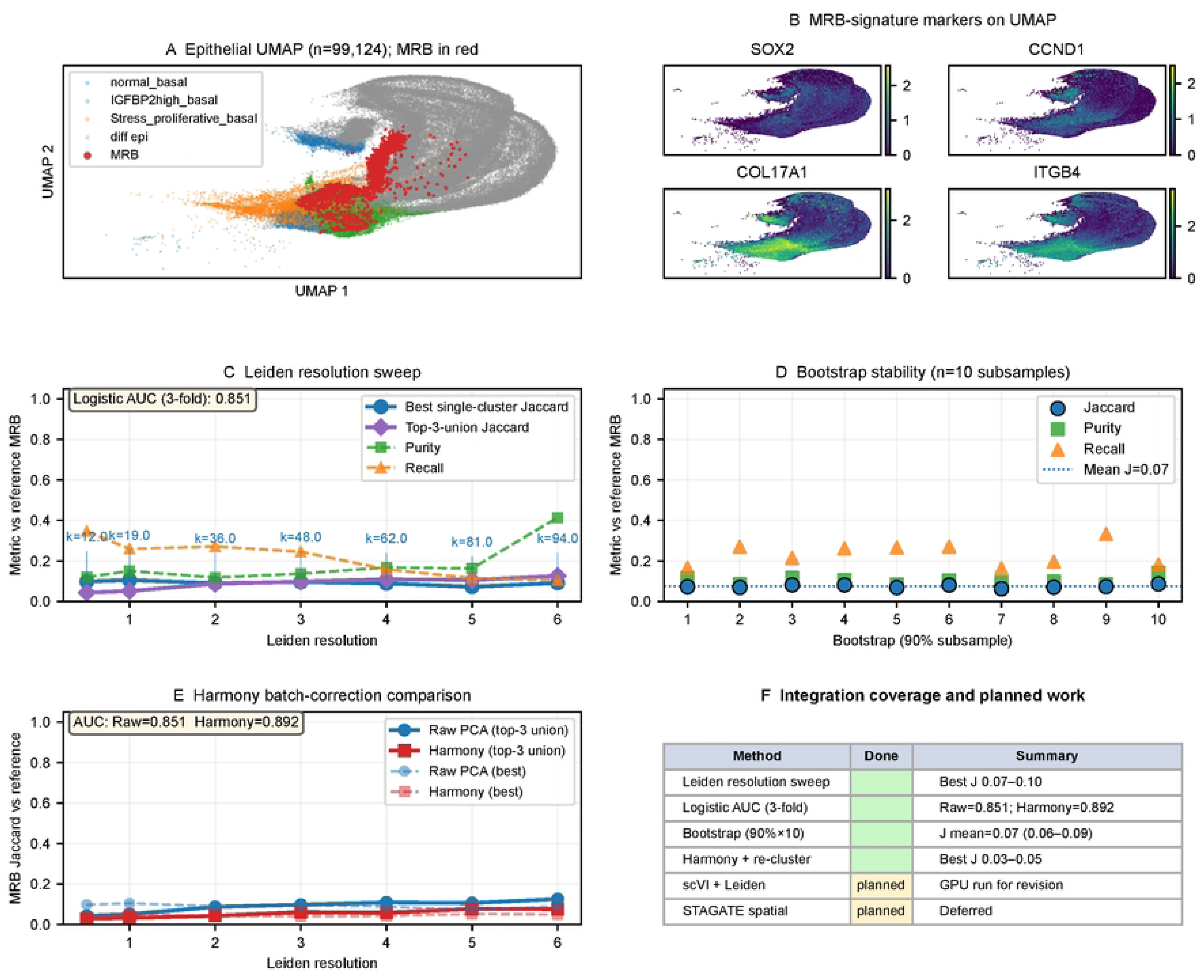

**S4_Fig.**
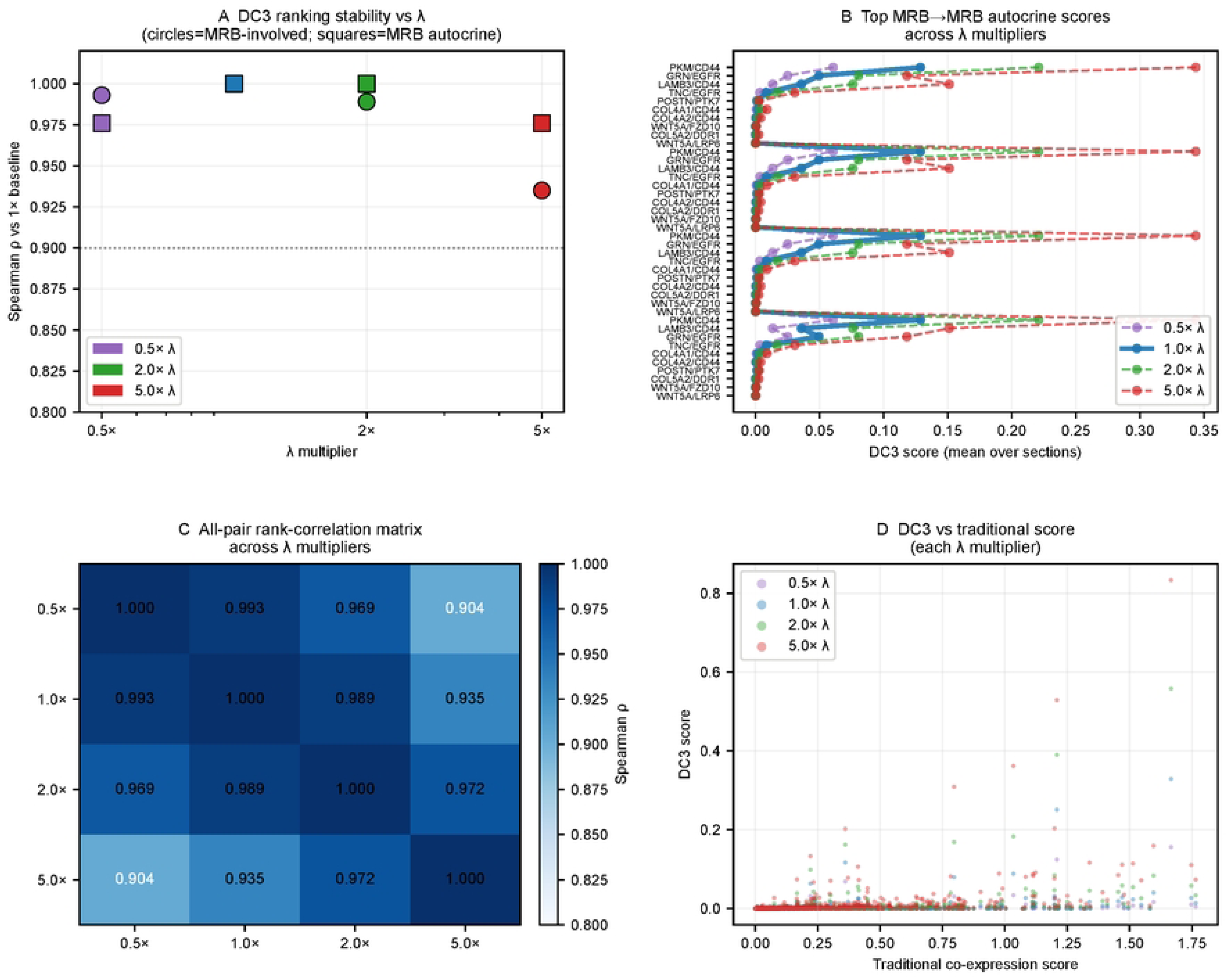

**S5_Fig.**
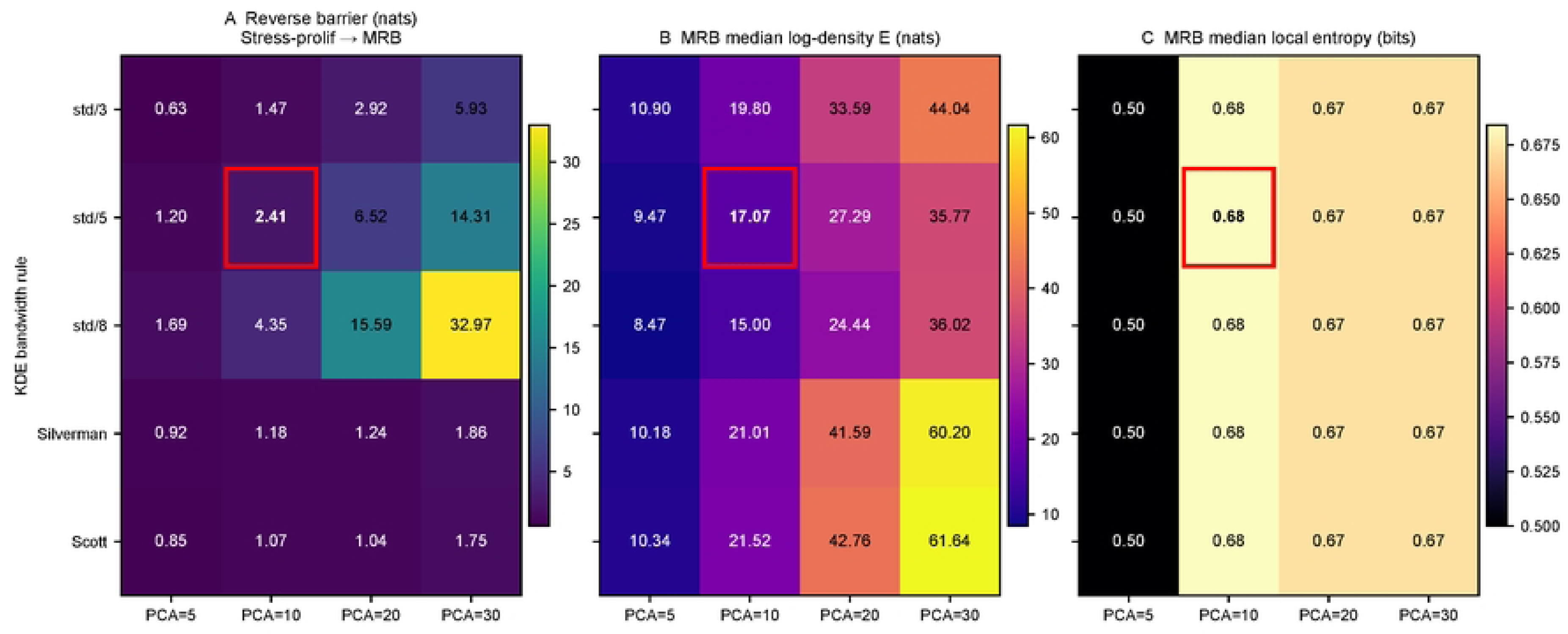

**S6_Fig.**
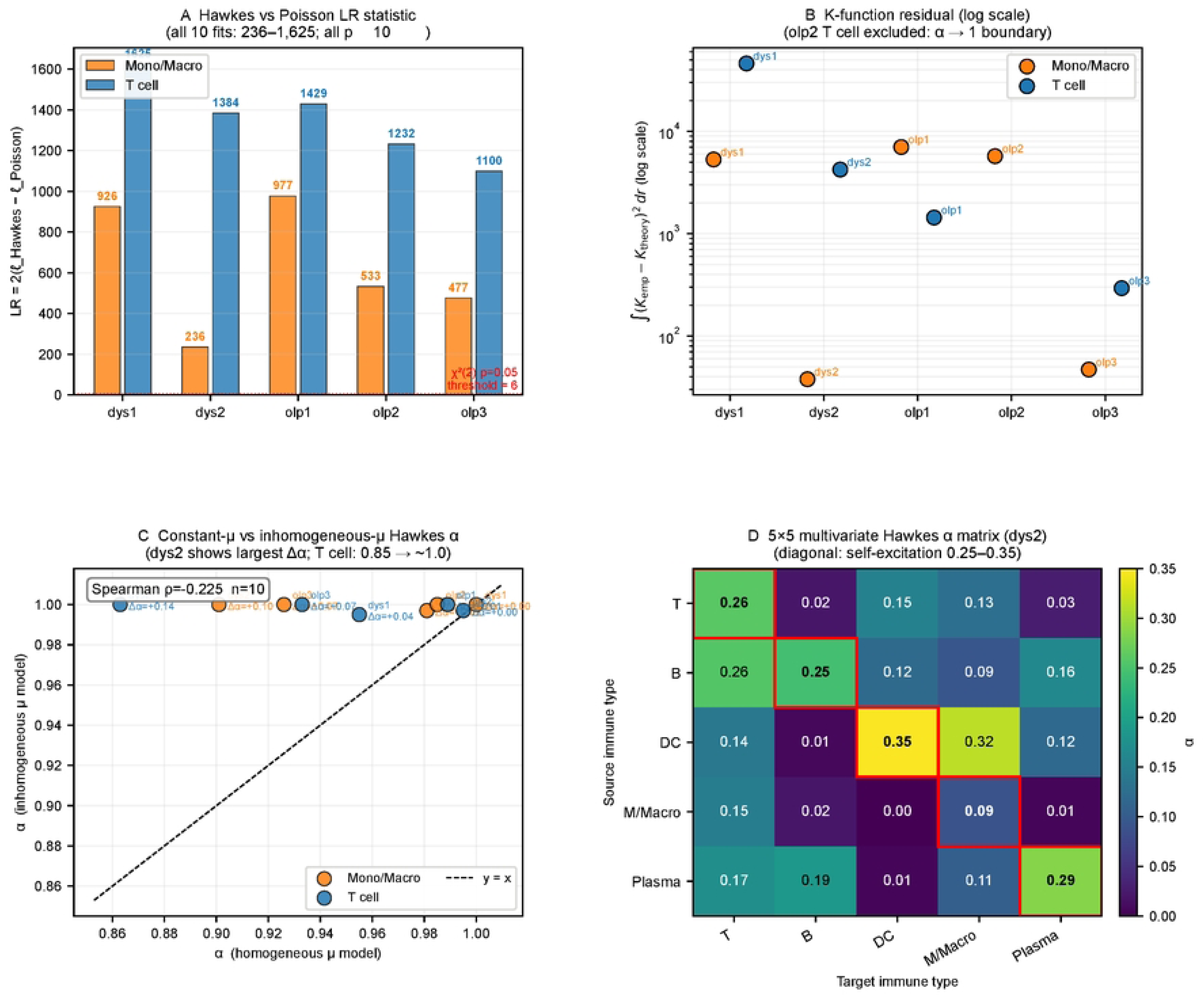

**S7_Fig.**
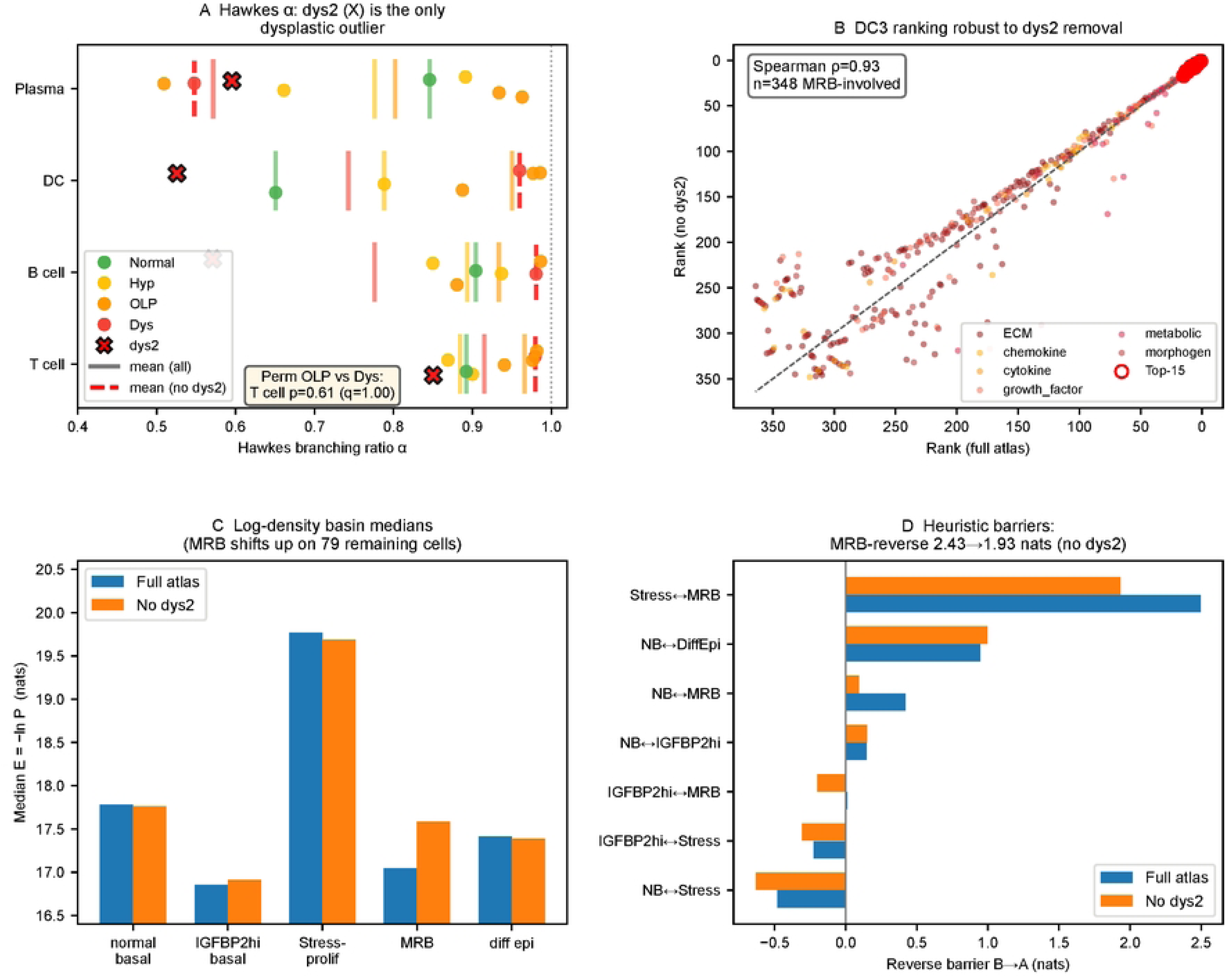

**S8_Fig.**
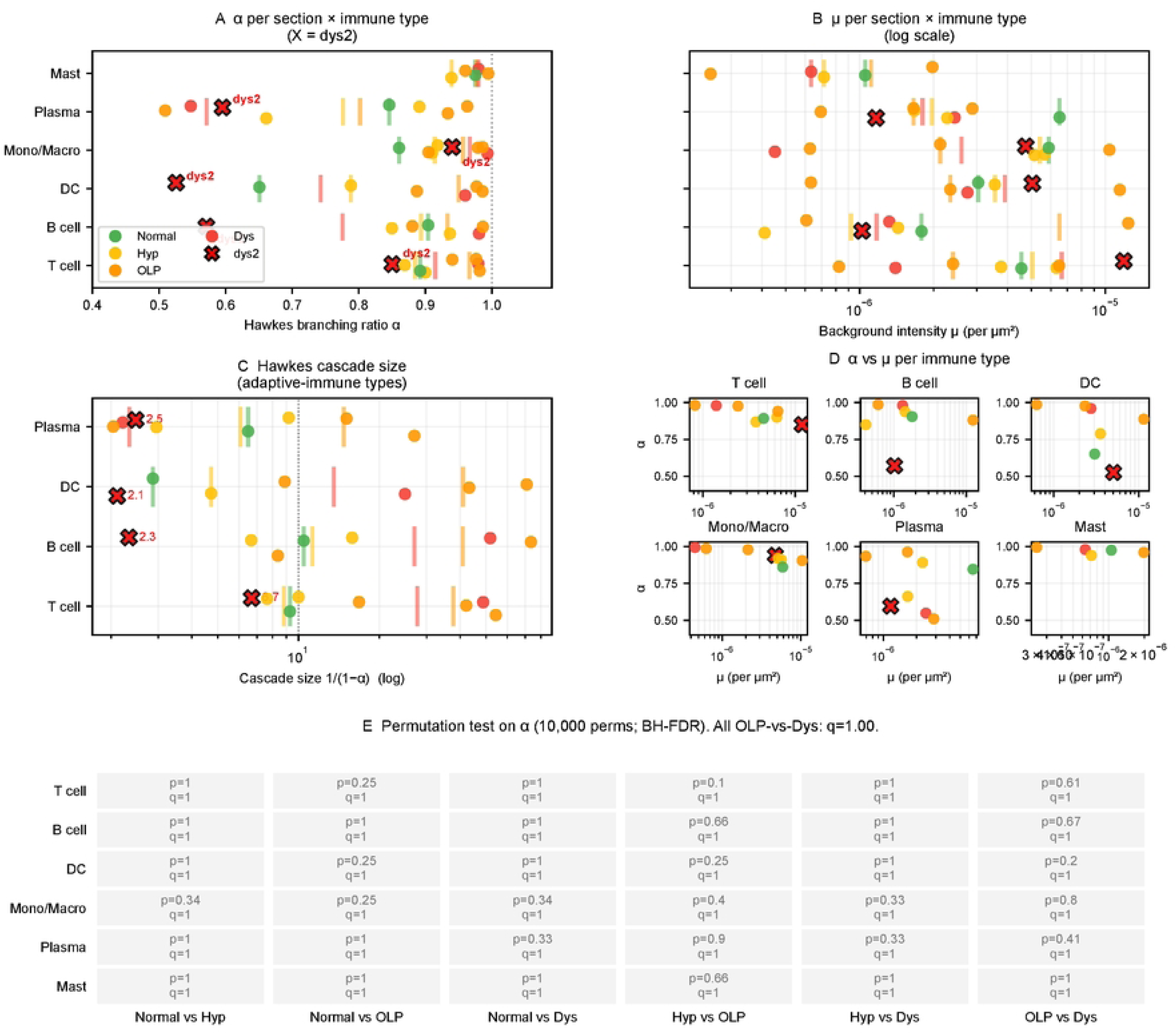

**S9_Fig.**
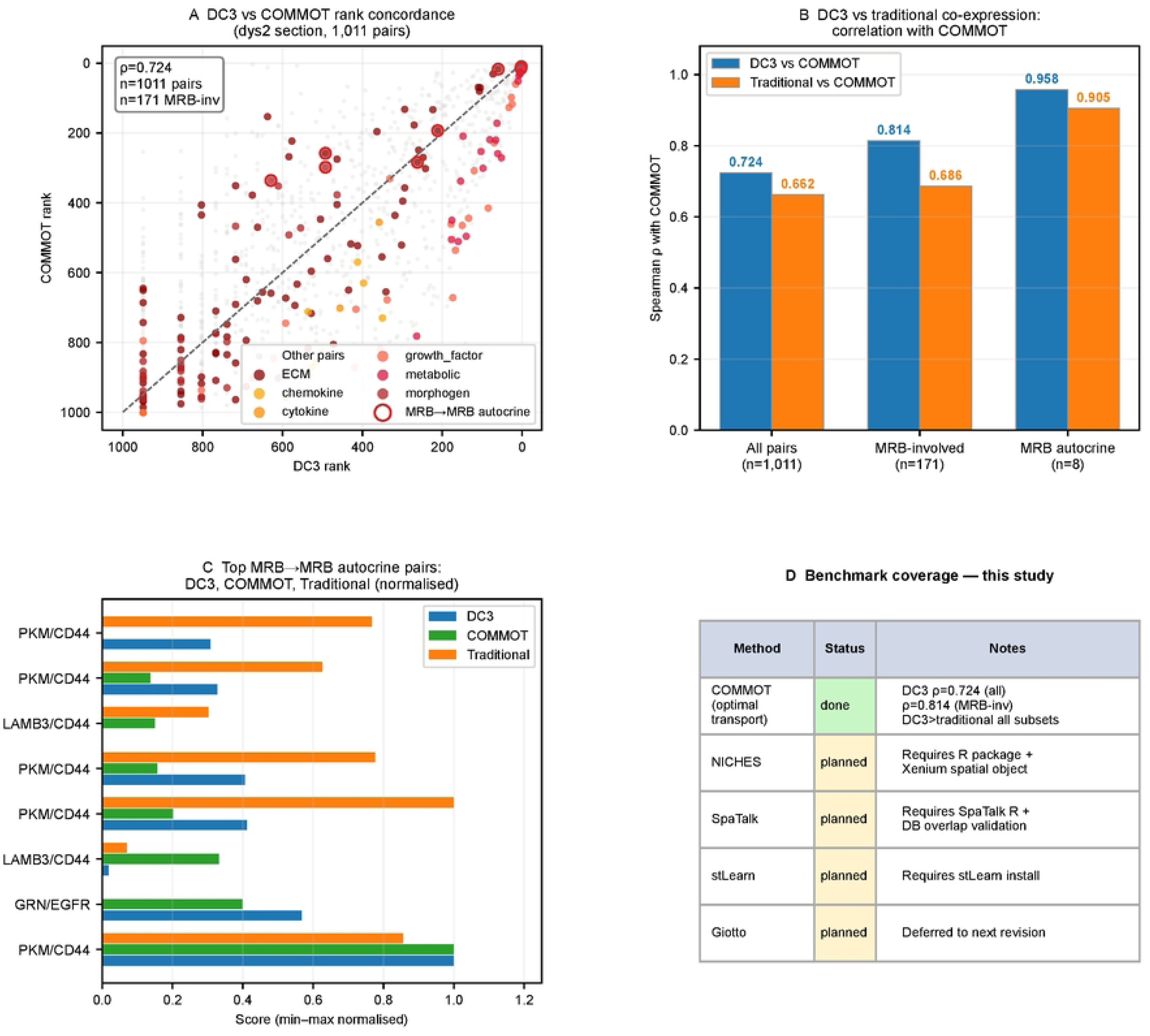

**S10_Fig.**
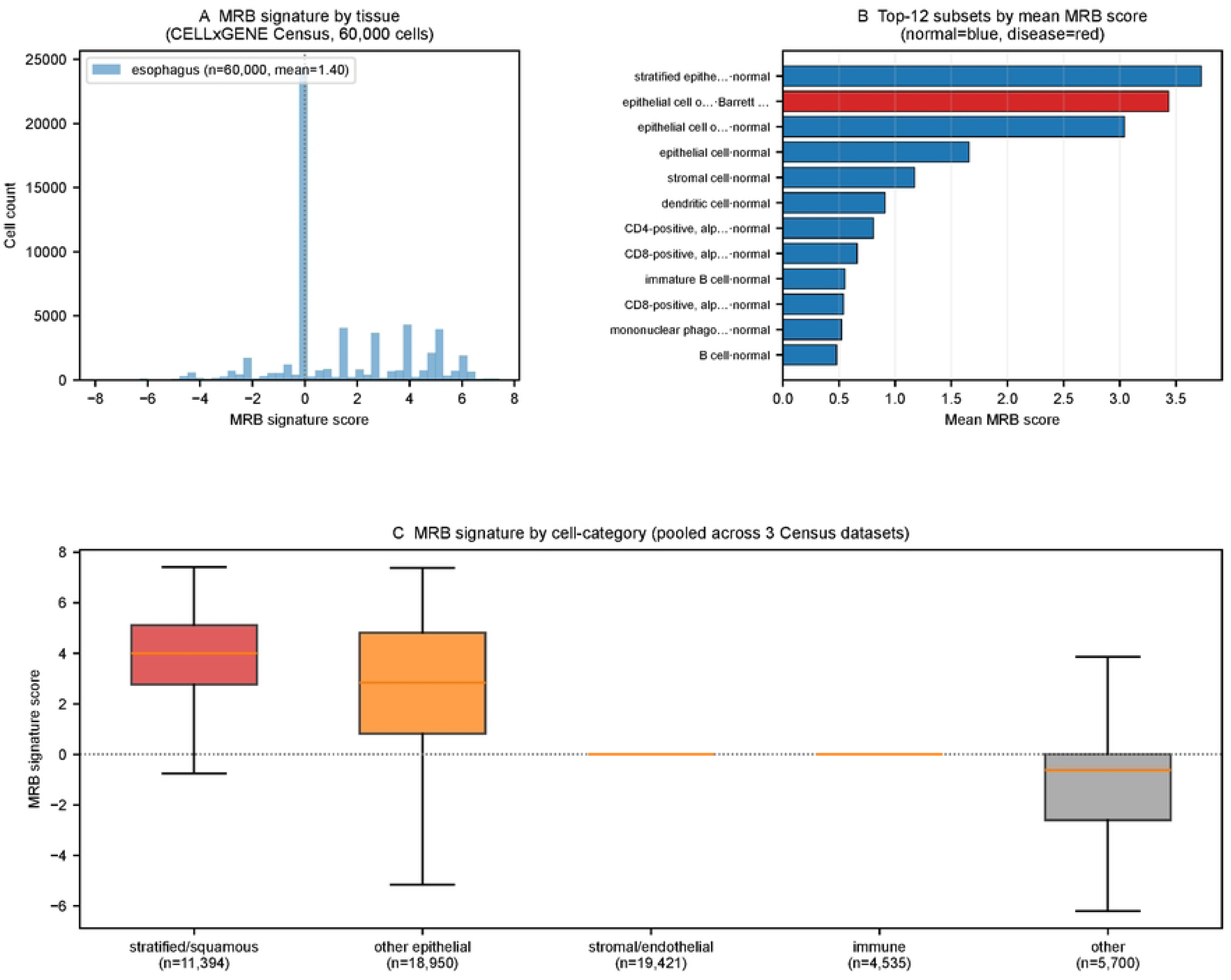

**S11_Fig.**
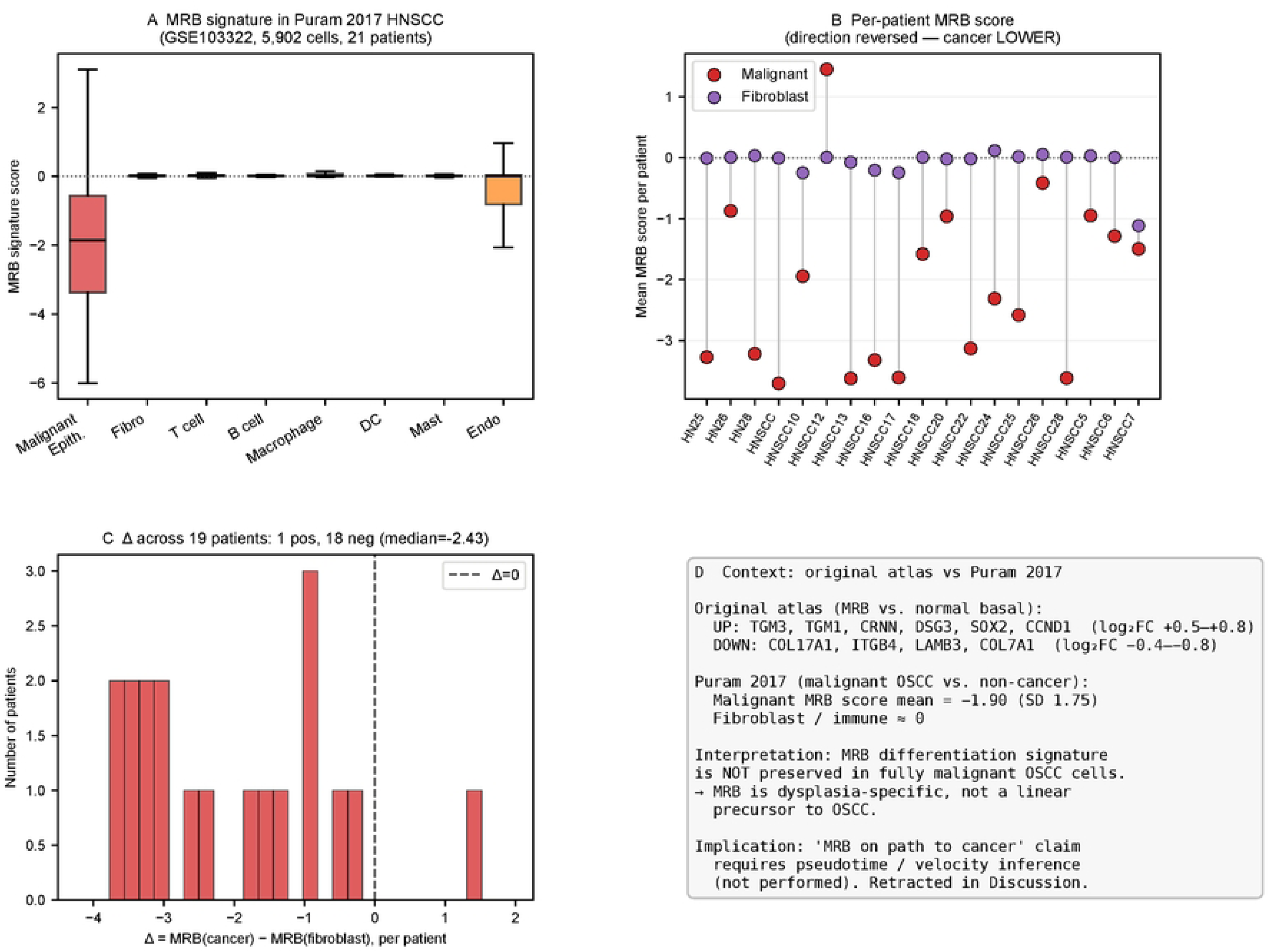

